# Chaparral wildfire shifts the functional potential for soil pyrogenic organic matter and nitrogen cycling

**DOI:** 10.1101/2024.08.18.607989

**Authors:** M. Fabiola Pulido Barriga, Amelia R. Nelson, Peter M. Homyak, Michael J. Wilkins, Sydney I. Glassman

**Author notes:** Corresponding authors: Sydney I Glassman, and M. Fabiola Pulido Barriga.

## Abstract

Wildfires reshape soil microbiomes and chemistry, enhancing nitrogen availability and leaving behind pyrogenic organic matter (PyOM), a difficult to degrade carbon substrate potentially used by pyrophilous or “fire-loving” microbes. Understanding whether pyrophilous bacteria can metabolize post-fire resources is critical for predicting the fate of both carbon and nitrogen. We explored how secondary succession of pyrophilous bacteria align with changes in functional gene composition, particularly genes related to PyOM degradation and microbial nitrogen metabolism using shotgun metagenomics on 30 burned and unburned soils collected at 17, 25, 34, 131, and 376 days after a high-severity wildfire in a fire-adapted chaparral in Southern California. In burned soils, genes for PyOM degradation increased over time by 167% and for inorganic nitrogen cycling by 117%, while unburned soils showed no significant changes. Genes encoding catechol and protocatechuate, intermediates in the PyOM degradation pathway, indicate that the easier-to-degrade ortho-cleavage pathways consistently dominated the burned plots. These pathways co-occurred with genes for nitrification and nitrogen retention, including assimilatory and dissimilatory nitrate reduction to ammonia (DNRA). We also identified 446 bacterial metagenome-assembled genomes (MAGs) and linked gene profiles to dominant taxa and found that increases in genes for PyOM and nitrogen cycling paralleled the dominance of pyrophilous *Massilia* and *Noviherbaspirillum*, which encoded distinct pathways for PyOM and inorganic nitrogen use over time. Together, these findings reveal previously unrecognized functional shifts in bacterial communities over a high-resolution successional timeline, providing insights into the long-term impact of fire on microbial-mediated ecosystem processes that shape soil carbon and nitrogen dynamics.

## 1.1 Introduction

Soil microbial communities support fundamental post-fire soil processes, including nitrogen (N) cycling through processes like immobilization, denitrification, and leaching (Turner et al., 2007), and carbon (C) cycling through sequestration and CO_2_ release during decomposition (Liu et al., 2023a; Raza et al., 2023). Despite evidence of rapid post-fire microbial succession (Pulido-Chavez et al., 2023), the functional dynamics of post-fire soil microbiomes remain poorly understood, largely due to the limited availability of high-resolution temporal sampling. As fire regimes increase in frequency and severity (Cunningham et al., 2024), understanding early functional changes in post-fire soil microbiomes is crucial for determining how microbes drive secondary succession and influence ecosystem functional recovery.

Wildfires have profound and lasting effects on global C and N cycling. Fires produce pyrogenic organic matter (PyOM)—a persistent C substrate formed during the incomplete combustion of vegetation (Knicker, 2011; Nave et al., 2011; Bird et al., 2015). PyOM’s complex chemical structure, with highly condensed aromatic and O-aromatic components (Chatterjee et al., 2012; De la Rosa et al., 2012; Chen et al., 2022), makes it difficult for microbes to degrade, typically breaking down over centuries (Singh et al., 2012), leading to long-term effects on post-fire nutrient dynamics (Haddock, 2010; Knicker, 2011; Bird et al., 2015). However, in-lab research suggests that PyOM breaks down into smaller structures within 10 months (Singh et al., 2014), which, while still containing aromatic components, may be more accessible to microbial enzymes, potentially facilitating microbial PyOM degradation. Additionally, wildfires alter soil chemistry by increasing pH, ammonium (NH□□), nitrate (NO□□), and urease activity (Ayiti and Babalola, 2022) which are intensified under high-severity fires that alter soil temperature and moisture (Neary et al., 1999; Moya et al., 2018), further impacting N cycling processes like nitrification and denitrification. However, wildfires typically reduce bacterial richness and shift community composition, favoring pyrophilous (“fire-loving”) bacteria, particularly from the Firmicutes and Actinobacteria phyla (Whitman et al., 2019; Enright et al., 2022; Pulido-Chavez et al., 2023). Yet, it remains unclear whether these bacteria possess the genes necessary to degrade PyOM and utilize the available post-fire N.

Pyrophilous bacteria can shape successional dynamics by altering resource use and driving predictable shifts in species composition (Pulido-Chavez et al., 2023) and metabolism (Luo et al., 2020; Nelson et al., 2022). They may drive secondary succession through a trait-based transition from thermotolerance to fast growth and ultimately PyOM utilization (Pulido-Chavez et al., 2023). In this context, early fast-growers metabolize labile C substrates, priming or preparing the soil for later-arriving microbes capable of degrading complex aromatic compounds in PyOM. This process likely relies on functional succession (Ni et al., 2023), where taxa integrate sequentially based on metabolic traits (Jackrel et al., 2019), potentially stabilizing microbial communities and enhancing ecosystem recovery. However, whether these functional changes occur simultaneously or influence microbial successional dynamics remains unclear.

Bacterial degradation of complex aromatic C substrates, such as lignin or PyOM occurs through the catechol and protocatechuate pathways, following either the meta- or ortho-cleavage pathway (Fuchs et al., 2011). The ortho-cleavage (β-ketoadipate) pathway is favored (Harwood and Parales, 1996; Parke et al., 2000; Buchan et al., 2004), due to its efficiency, higher energy yield, and enzyme specificity (Harwood and Parales, 1996; Bugg et al., 2011; Fuchs et al., 2011), which may be particularly relevant in post-fire landscapes (20). For example, *Massilia*, a well-known pyrophilous genus in the Burkholderiales (Enright et al., 2022; Pulido-Chavez et al., 2023), may degrade both lignin (Wang L, 2016) and PyOM (Pérez-Pantoja et al., 2012) via the ortho-cleavage pathway (Pérez-Pantoja et al., 2012). Beyond aromatic compounds, wildfires also increase short-chain alkanes (González-Pérez et al., 2004; Badía et al., 2014), such as propane, a relatively more labile C source preferentially consumed by microbes (Moucawi et al., 1981; Abbasian et al., 2015; Thomas et al., 2021). However, it remains undocumented how and when pyrophilous bacteria metabolize these different C sources or how their C substrate preference shifts during post-fire microbial succession.

Microbial metabolism also plays a crucial role in post-fire N cycling, influencing nutrient availability and ecosystem recovery. While post-fire greenhouse gas (GHG) emissions are common (Stephens and Homyak, 2023), microbial communities can mitigate these emissions by converting abundant NH□□ and NO□□ into microbial biomass via assimilatory and dissimilatory nitrate reduction (DNRA) (Neary et al., 1999). This process may shape microbial succession by favoring taxa with NO□□ assimilation genes. Indeed, rapid post-fire shifts in microbial composition (Pulido-Chavez et al., 2023) likely reflect changes in microbial metabolism, influencing the recovery of inorganic N, such as NH□□ and NO□□ which undergo rapid changes in the first post-fire years (Stephens and Homyak, 2023). Notably, pyrophilous bacteria analyzed 3–10 years post-fire retain substantial N-cycling and PyOM degradation genes (Dove et al., 2022; Nelson et al., 2022; Whitman et al., 2022), indicating their adaptation to shifting soil nutrient conditions. This suggests that microbial communities adapt to shifting post-fire conditions and may influence ecosystem recovery through functional changes that drive microbial succession.

Here, we used shotgun metagenomics to examine how pyrophilous bacteria drive post-fire microbial functional shifts in C and N cycling, leveraging soils where we previously characterized microbiomes across nine time points from 2 weeks to 1 year post-fire (Pulido-Chavez et al., 2023). Focusing on five key time points corresponding to major bacterial turnover events (Pulido-Chavez et al., 2023), we examined whether bacterial secondary succession aligns with shifts in functional gene composition, particularly genes related to PyOM degradation and microbial N metabolism. We used metagenomically assembled genomes (MAGs) to link functional changes to dominant pyrophilous bacteria over time. We tested the hypotheses that H1) The relative abundance of PyOM degradation genes will increase over time as alkane degradation genes decrease, reflecting microbial adaptation to PyOM as a C source; H2) Genes involved in microbial N metabolism will increase over time, indicating shifts in bacterial functional potential in response to post-fire N availability; and H3) Shifts in gene abundances will correspond with bacterial community shifts, particularly the increasing dominance of pyrophilous bacteria over time.

### 1.2 Methods

#### 1.2.1 Sampling

In August-September 2018, the Holy Fire burned 94 km² of Manzanita-dominated chaparral in Southern California. Using BAER soil burn severity maps (<u>USGS</u>), we established nine plots (six burned, three unburned) 17 days post-fire, selecting sites with similar pre-fire vegetation, elevation, and aspect. These plots spanned Entisols (Typic Xerorthents) mapped in the Cieneba series and Mollisols (Lithic Haploxerolls) in the Friant series (Pulido-Chavez et al., 2023). Each plot contained four 1 m² subplots in each cardinal direction for temporal soil sampling of the top 10 cm of mineral soil beneath the ash or duff layer (Fig. S1). To assess soil burn severity at the plot level, we used ash depth as a proxy, averaging three measurements per subplot. Additional details of the experimental design can be found in Pulido et al. (Pulido-Chavez et al., 2022). Soil samples were collected at nine time points ranging from 2 weeks to 1 year post-fire and stored at −80°C within 24h of collection (Pulido-Chavez et al., 2023). DNA was extracted from 0.25g of 2mm sieved soil and we performed 16S rRNA and ITS2 Illumina MiSeq sequencing of all samples (Pulido-Chavez et al., 2023). Building on our previous study of successional dynamics, we selected the five time points with the largest bacterial turnover events (17, 25, 34, 131, and 376 days post-fire) and sufficient high-quality DNA for metagenomic sequencing (Pulido-Chavez et al., 2023).

#### 1.2.2 Metagenomic sequencing

From each of the five timepoints, we selected 3 burned and 3 unburned subplots, totaling 30 metagenomes. Samples with DNA concentrations of 2.5 ng/µl and 230/260 above 1.8 were selected for metagenomes. Samples were collected from unburned plot 8 and two closely situated burned plots with similar microbial richness and biomass (plots 2 and 3; Fig. S1a). Metagenomic library preparation and sequencing were performed at the Department of Energy Joint Genome Institute (JGI) for 17 samples and at the University of California, Irvine (UCI) Genomics High-Throughput Facility for 13 samples. Libraries were prepared using the Kapa Biosystems library preparation kit at JGI and the Illumina Nextera Flex library preparation kit at UCI. Sequencing was performed on the Illumina NovaSeq 6000 platform with 151 bp paired-end reads and a depth of 25 Gb/sample.

#### 1.2.3 Metagenomics binning

Sequence adapters were removed from the raw reads using BBduk-v38.89 (<u>sourceforge.net/projects/bbmap/</u>) with the following parameters (ktrim=r, k=23, mink=11, hdist=1). Raw reads were trimmed with Sickle – v1.33 (Joshi NA, 2011), and the quality of trimmed reads was assessed with FastQC – v0.11.2 (Andrews, 2010). Trimmed reads were assembled into contiguous sequences (contigs) with the de novo de Bruijn assembler MEGAHIT – v1.2.9 (Li et al., 2014) with a min kmer of 27, max of 127, and increment of 10. Per-contig coverage per sample was calculated using CoverM contig v0.6.0 (Woodcroft, 2007) with the trimmed mean method—i.e., the average number of aligned reads per position after removing the most deeply and shallowly covered regions—while retaining only mappings with >95% identity and a minimum alignment length of 75% (Parks et al., 2015). Assembled contigs (>2,500 bp) were binned using MetaBAT2 v2.12.1 with default parameters (Kang et al., 2015) to assemble metagenome-assembled genomes (MAGs). MAG quality was estimated using checkM v1.1.2 (Parks et al., 2015) and taxonomy was assigned using GTDB-Tk v2.1.1 (Chaumeil et al., 2019). To create the final medium and high-quality MAG database, low-quality MAGs (<50% completion, >10% contamination (Bowers et al., 2017) were removed and the dataset was dereplicated using dRep v3.0.0 (Olm et al., 2017).

#### 1.2.3 Contig annotation

We assessed microbial preferences for inorganic versus organic N cycling, focusing on urea, the largest contributor to the organic N cycle through hydrolysis into NH□□ (Fernández-García et al., 2020). For C cycling, we focused on short-chain alkanes as a more readily degradable C source and PyOM genes, focusing on catechol and protocatechuate, key intermediates of aerobic aromatic degradation (Fuchs et al., 2011), as the more complex C sources.

To improve retention and to comprehensively annotate metagenomes, we annotated all assembled contigs using three different databases. First, contigs were annotated using HMMER (Eddy, 2011), with hidden Markov models (HMMs) against Kofamscan HMMs (Aramaki et al., 2020) to identify specific genes according to previously established methods (Nelson et al., 2022). Second, we used the curated Calgary approach to AnnoTating HYDrocarbon degradation genes (CANT-HYD) database, which uses HMMs for annotating marker genes involved in aerobic and anaerobic hydrocarbon degradation pathways (Khot et al., 2022). Third, we used DRAM (Shaffer et al., 2020), an annotation tool to profile our contigs through multiple databases including KEGG (Kanehisa and Goto, 2000), UniRef90 (Suzek et al., 2007), MEROPS (Rawlings et al., 2014), and Mmseqs2 (Steinegger and Söding, 2017) with the best hits, based on a bit score of 60, reported for each database. Together, these methods improve annotation accuracy by cross-validating results, capturing both broad metabolic functions and specialized enzymatic activities, and improving confidence in gene function assignments. Contigs counts across samples were normalized using the contig length corrected trimmed mean of M values (geTMM) (Smid et al., 2018) using EdgeR (Robinson et al., 2010), adjusting for library depth and gene length (Capo et al., 2021; Leleiwi et al., 2023; Santos-Medellin et al., 2023). For contigs with duplicate annotations, we manually curated to select the annotation with the lowest e-value as the best functional annotation.

We identified genes of interest using DESeq analysis (details below) and linked them to our MAG catalog via NCBI BLAST (Bethesda, 2008), filtering to include only hits with 100% identity and a bit score ≥ 100, which we used to identify the specific taxa that encoded key genes related to PyOM, alkane, and organic and inorganic N cycling pathways. See Supplementary Data 1 for additional information regarding metagenomic sequencing, assemblies, and annotations, and see Fig. S2 for an overview of bioinformatic procedures employed.

#### 1.2.4 Statistical analysis

All statistical analyses and figures were done in R version 4.2.3 (RCoreTeam, 2023). To determine how fire, time, and their interaction impacted the C and N gene composition, we employed Permutational Multivariate Analysis of Variance (PERMANOVA;(Anderson, 2017)) via adonis2 in the vegan package (Oksanen et al., 2022) with 9999 permutations with plot as random effect and visualized with nonmetric multidimensional scaling (NMDS). Ordinations were based on Bray–Curtis distance matrixes using the ordinate function in the phyloseq package (McMurdie and Holmes, 2013).

To determine if bacterial functional changes mirrored previously observed rapid community compositional changes (Pulido-Chavez et al., 2023), we plotted the most abundant bacterial genera identified with 16S amplicons at each post-fire timepoint. We also calculated the homogeneity of variance using permutest with 9999 permutations and plot as random effect to determine which time points differed for burned and unburned pathways independently. Principal coordinates analysis (PCoA) was used to visualize the differences in the homogeneity of variance of each functional pathway employing Bray-Curtis dissimilarities in the vegan package.

We performed generalized negative binomial regressions on burned and unburned plots for each pathway to test how the number of PyOM, alkane, inorganic and organic N cycling genes changed over time. Nestedness level was assessed using null model and Akaike criteria. For burned plots, models for PyOM, alkane and organic N included plot as a random effect, while unburned plots included subplot. In contrast, inorganic N models for burned plots included plot, subplot and time as random effects, while unburned models for both organic and inorganic N included time and subplot as random effects. Time was scaled and centered for all models. We visualized changes in microbial functional potential over time by plotting the sum geTMM for each pathway of interest at each time point, separately for burned and unburned plots, using phyloseq version 1.42.0 (McMurdie and Holmes, 2013).

Lastly, to identify the PyOM, alkane, inorganic and organic N genes that were differentially abundant in burned versus unburned plots at each sampling time point, we employed DESeq2 v1.38.3 (Love et al., 2014). To evaluate the response of functional gene potential over time, we created separate DESeq2 models for each pathway at each time point, determining significance with the Wald test and an adjusted p-value of < 0.05 and an absolute log2 fold-change > 0. The total count of differentially abundant genes was visualized as a bar plot for each pathway over time. We used the differentially abundant genes from the burned communities to create metabolic maps for PyOM and inorganic N degradation pathways at each time point. Genes identified as abundant by DESeq at each time point were linked to their respective taxonomy using our MAG dataset via BLAST (Bethesda, 2008). The taxonomy and functional annotation of the selected genes were visualized using ggplot2. All R scripts are publicly available on GitHub: https://github.com/pulidofabs/Chaparral-Metagenomes.

### 1.3 Results

Metagenomic sequencing of 30 burned and unburned chaparral soils generated 1.2 Tb of data and 2,266,623 annotated genes relevant to post-fire microbial metabolisms. The significantly abundant genes, as determined by the DESeq results, were linked to 17 of the 446 MAGs identified from this dataset (see Supplementary Data 1C for additional details on MAG quality and taxonomy). MAGs include the pyrophilous bacterial genera *Massilia* and *Noviherbaspirillium* that previously dominated our ASV dataset (Pulido-Chavez et al., 2023).

#### 1.3.1 Wildfire altered the profile of functional genes for PyOM and inorganic N cycling

Fire and time significantly impacted abundances of functional genes associated with degradation of PyOM (Fig. 1a), alkanes (Fig. 1b), and cycling of inorganic (Fig. 1c) and organic N (Fig. 1d). Further, a significant fire-by-time interaction affected degradation of PyOM and cycling of inorganic and organic N, such that genes changed over time in burned but not unburned plots (Table S1). Fire was the greatest driver of differences in functional gene profiles, accounting for 15% to 16% of the variation, with time and fire-by-time interactions accounting for 5-6% of the variance in C and N gene functional profiles (Table S1).

**Figure 1.**
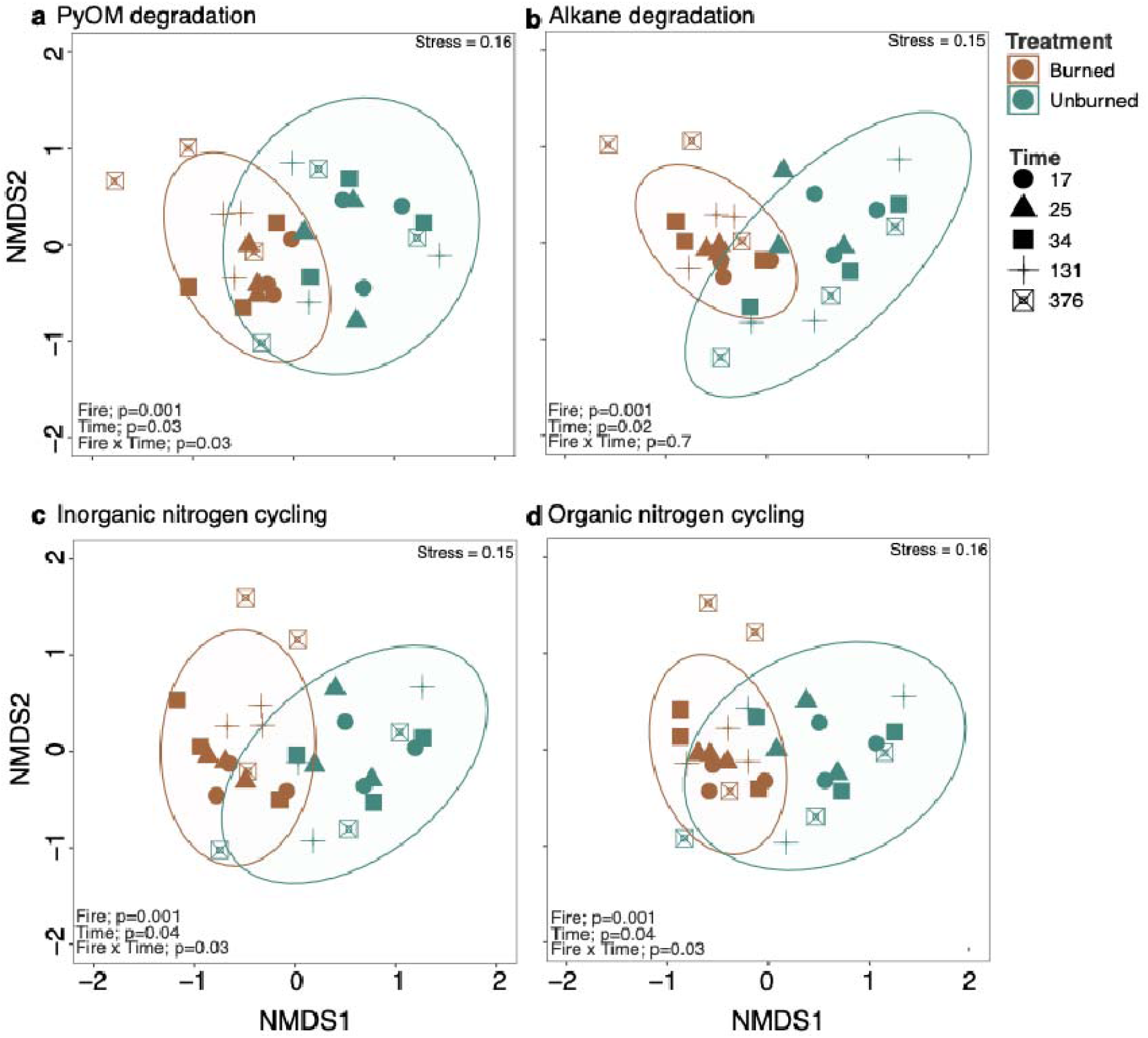
Non-metric multidimensional scaling (NMDS) of geTMM normalized a) PyOM b) alkane c) inorganic nitrogen and d) organic nitrogen cycling genes within the microbial communities of burned and unburned soil samples (color) and time since fire (days; shape). Fire and time since fire effects and stress based on PERMANOVA (p <0.05). Ellipse represents the 95% confidence level.

While unburned plots showed no turnover in functional gene profiles for PyOM (Fig. 2a), or alkane degradation (Fig. 2c), or inorganic (Fig. 2e) and organic N cycling genes (Fig. 2g), burned plots exhibited multiple functional turnover events for PyOM (Fig. 2b), alkane (Fig. 2d), inorganic (Fig. 2f), and organic N genes (Fig. 2h; Table S2). Specifically, PyOM degradation and inorganic N cycling genes turned over at 34 and 131 days, alkane degradation genes at 25 and 131 days, and organic N processing genes at 25, 34, and 131 compared to 17 days post-fire (Fig. 2).

**Figure 2.**
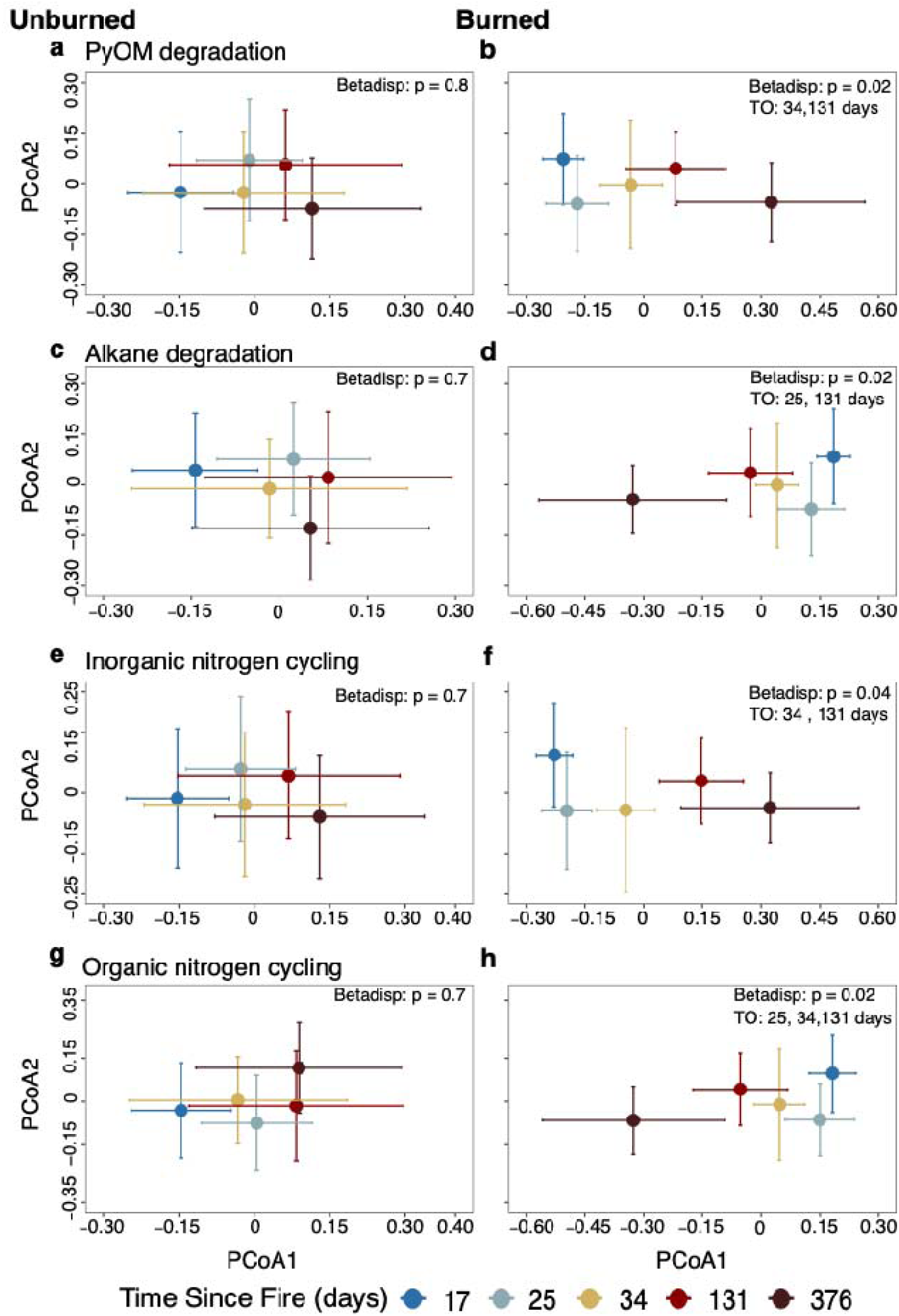
Principal component analysis (PCoA) of the mean and standard error Bray-Curtis community dissimilarity at each timepoint for the unburned and burned a,b) PyOM; c,d) alkane; e, f) inorganic nitrogen and g, h) organic N (urea) functional potentials. Note that timepoints further apart with nonoverlapping standard error bars indicate a community turnover, whereas overlapping standard error bars represent a lack of compositional turnover, with the first sampling time point at 17 days representing the base level. Thus, the first turnover was initiated at 25 days. There were no significant turnover (TO) dates for any gene functional potentials in the unburned plots. The days with significant turnovers in the burned plots are listed in the upper right corners. Significance based on betadisper, which measures the dispersion/variability of the data within groups to measure community turnover.

#### 1.3.2 Time differentially affected PyOM cycling pathways

A significant fire-by-time interaction led to a 167% increase in total geTMM-normalized gene abundance for PyOM degradation in burned plots from 17 to 376 days post-fire (Fig. 3b), while alkane cycling genes increased by 91% (Fig. 3d). In contrast, PyOM and alkane gene abundances remained relatively stable in unburned plots (Fig. 3a; Table S3). This time-dependent increase was also observed at the individual pathway level, where PyOM degradation pathways, including catechol, protocatechuate, and β-ketoadipate degradation (Fig. S3; Table S5), and the propane pathway for alkane degradation increased over time in burned but not unburned plots (Fig. S4; Table S6). This pattern likely reflects post-fire bacterial diversity recovery (Fig. S5b), which increases in burned plots over time as does the total number of PyOM degradation genes (Fig. S6).

**Figure 3.**
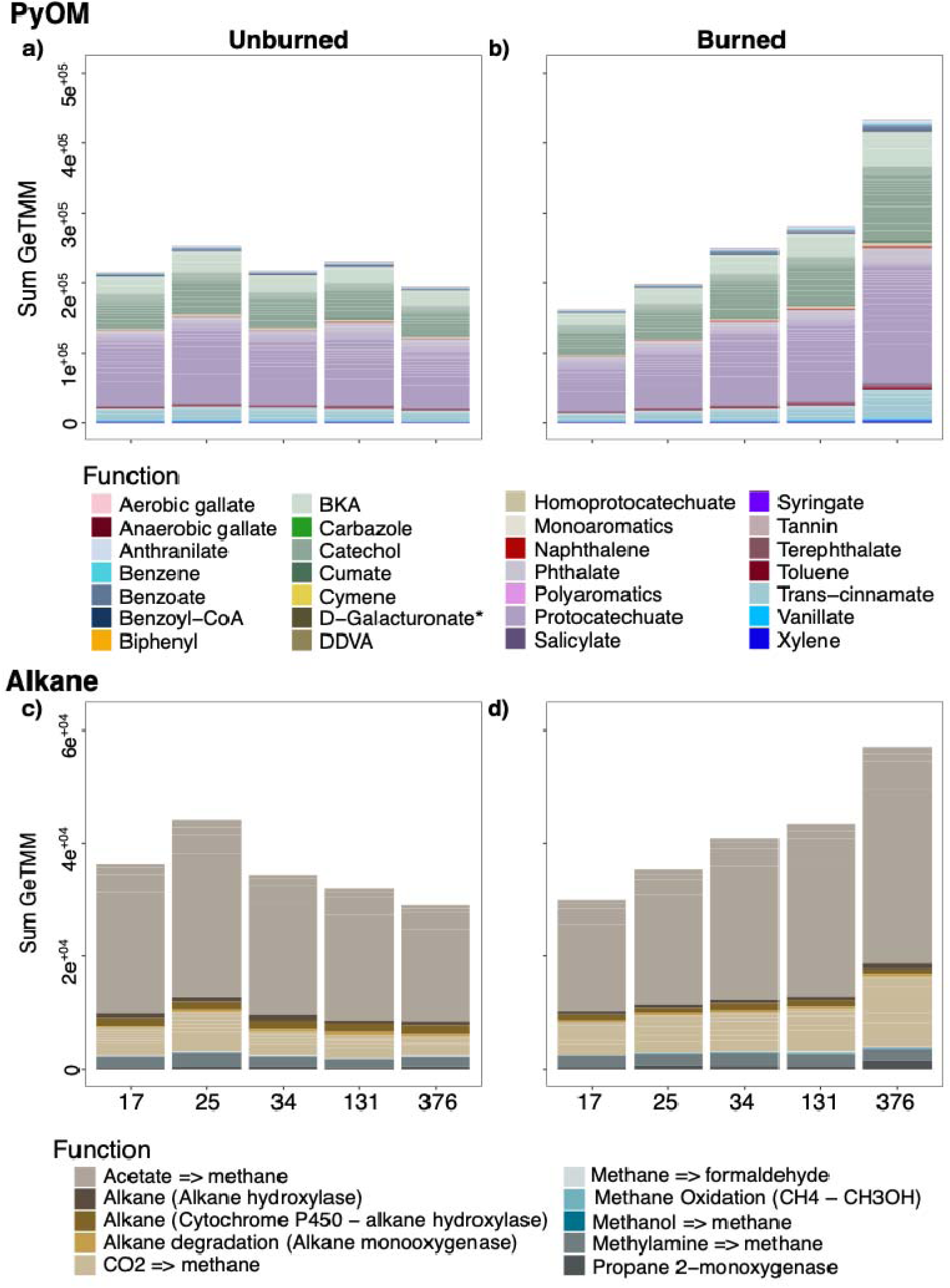
Summed geTMM-normalized gene abundance over time for PyOM degradation in (a) unburned and (b) burned plots, and alkane degradation in (c) unburned and (d) burned plots. Colors represent gene function. Generalized Negative Binomial Models indicate a significant increase in relative abundance over time in burned plots, while unburned plots remained stable (Table S1).

#### 1.3.3 Time differentially affected N cycling pathways

A significant fire-by-time interaction led to a 117% increase in total geTMM-normalized gene abundance for inorganic N cycling in burned plots from 17 to 376 days post-fire, while organic N cycling genes increased by 108% (Fig. 4; Tables S3, S4). In contrast, both inorganic and organic N gene abundances remained relatively stable in unburned plots. This temporal increase was also evident at the pathway level, with nitrification, DNRA, and NO□□ assimilation increasing in burned plots (Fig. S7; Table S7). This pattern may also be linked to bacterial diversity recovery (Fig. S4d), which increases in burned plots over time as does the total number of inorganic N cycling genes (Fig. S8). Interestingly, denitrification genes (NO□□ → NO□□ → NO; *narGZ, nrxA, nirK*) significantly decreased over time in unburned plots but remained stable in burned plots (Fig. S7; Table S7). Organic N cycling pathways did not differ between burned and unburned plots over time (Fig. S9; Table S8), though two pathways increased in abundance overtime in burned plots (Fig. S9).

**Figure 4.**
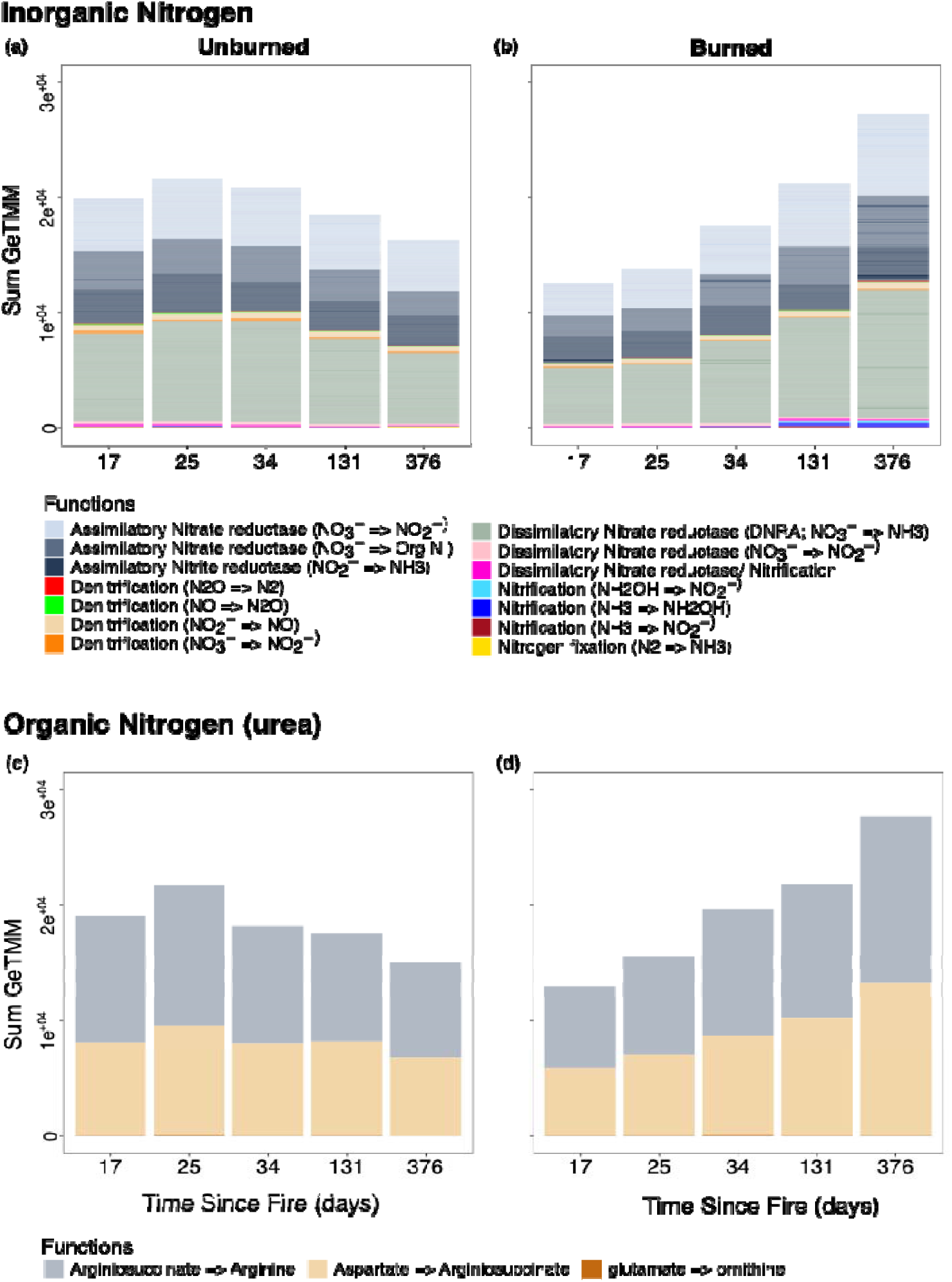
Summed geTMM-normalized gene abundance over time for inorganic N cycling genes in (a) unburned and (b) burned plots, and organic N cycling in (c) unburned and (d) burned plots. Colors represent gene function. Generalized Negative Binomial Models indicate a significant increase in relative abundance over time in burned plots, while unburned plots remained stable (Table S1).

#### 1.3.4 Wildfire selects for ortho-cleavage aromatic degradation pathways

Wildfire resulted in more differentially abundant PyOM degradation genes in burned compared to unburned soils (692 compared to 258; p_adj_ <0.05; Fig. 5a), with significant temporal variation in burned plots (p_adj_ < 0.05; Figs. 5a, 6). Ortho-cleavage pathways, including protocatechuate, catechol and β-ketoadipate, were differentially abundant and dominated burned plots from 17 to 131 days post-fire (p_adj_ <0.05; Fig. 5a, 6). In contrast, the meta-cleavage pathway, specifically protocatechuate meta-cleavage, was only dominant at 17 days post-fire (p_adj_ <0.05; Fig. 5a, 6). Pathways involving phthalate, toluene, and trans-cinnamate, which feed into the catechol or protocatechuate pathways, were also differentially abundant in burned plots between 17 and 131 days post-fire (p_adj_ < 0.05; Fig. 5a, 6). Conversely, unburned plots were enriched in lignin degradation genes (e.g., *ligABCIJKM* and *galC*) at 17 and 25 days (Fig. 5a, 6). Additionally, alkane degradation genes for acetate (*ACSS1_2, acs*, and *pta*) were present at all post-fire time points (Fig. 5b), while genes for propane 2-monooxygenase, alkane hydroxylase, and alkane monooxygenase were overrepresented only in burned plots at 17 days post-fire (p_adj_ < 0.05; Fig. 5b).

**Figure 5.**
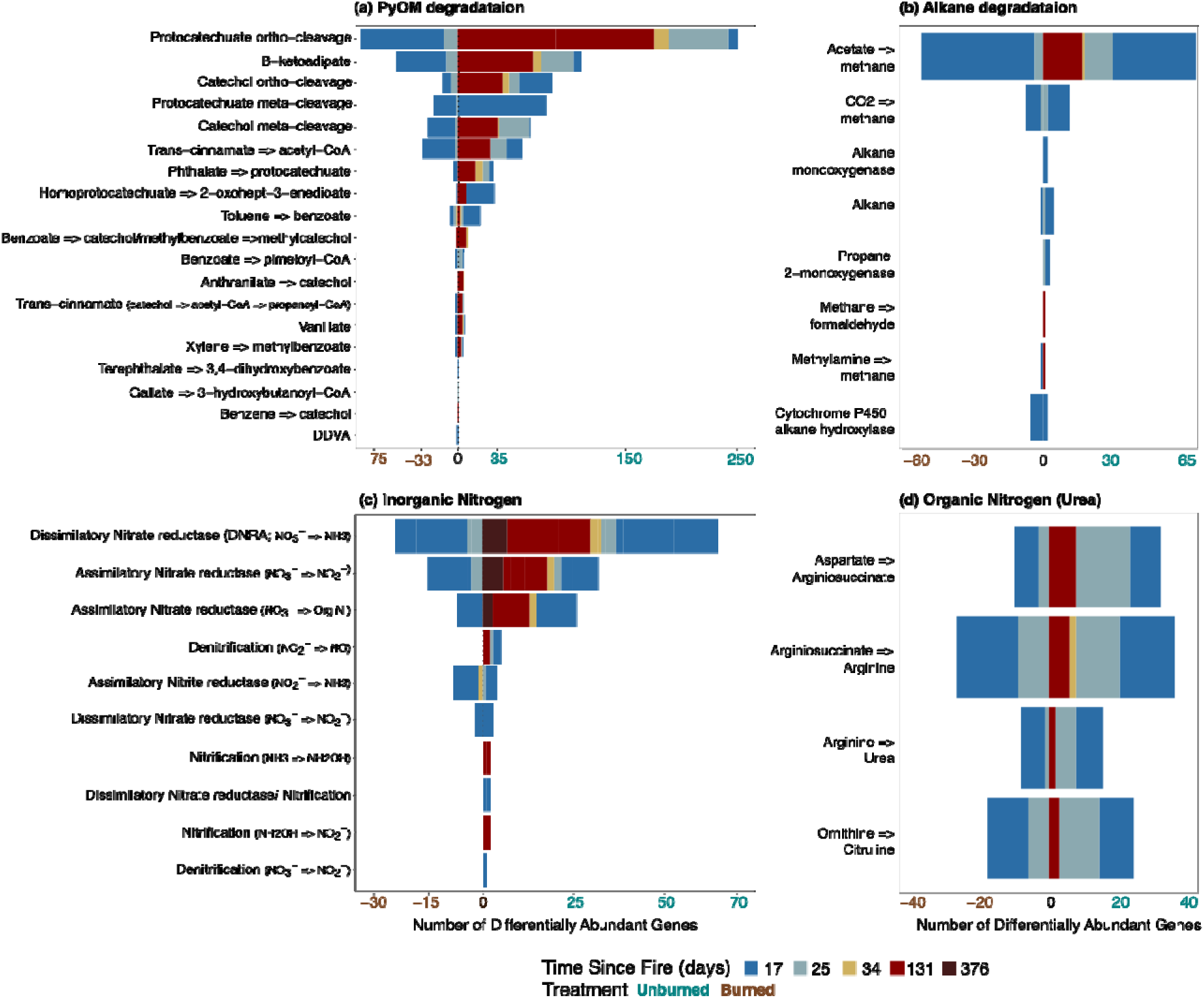
Total number of genes that were differentially abundant in the burned plots (brown) versus the unburned plots (blue green; x axis) per each metabolic pathway, colored by each sampling time point for a) PyOM, b) alkane, c) inorganic nitrogen and d) organic N (urea) cycling. Significance based on DESeq2 Wald test (adjusted P value < 0.05 and absolute log2 fold-change >0).

**Figure 6.**
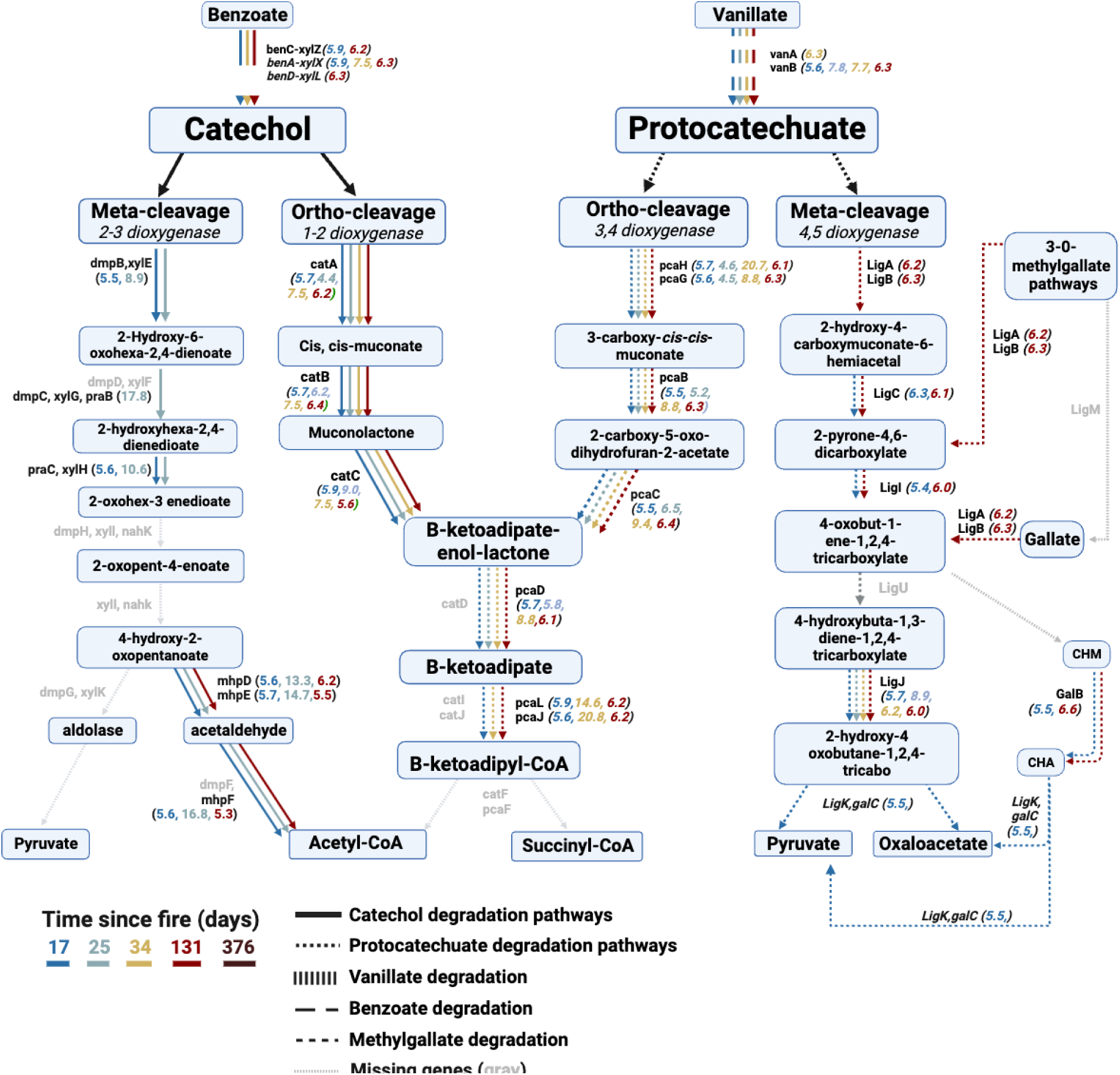
The catechol and protocatechuate degradation pathways with arrows indicating genes that were differentially expressed in the burned treatment at each time point (17, 25, 34, 131 and 376 days post-fire; Wald’s test in DESeq2; *p*<0.05). Line type represents catechol (solid) and protocatechuate (dashed) cycling pathways and gray solid lines represent pathways that were not expressed during post-fire year one. Numbers in parentheses represent the mean log2fold change (DESeq2) for each represented gene and are colored by the timepoint in which that gene was significantly abundantly represented.

#### 1.3.5 Wildfire increased genes for N retention

Wildfire increased the number of differentially abundant N cycling genes in burned compared to unburned soils (143 vs. 55; Fig. 5c, 7). At all timepoints, genes involved in NO_3_^-^ retention pathways, including assimilatory (*nasABC, NRT, narK, nrtP*, *nirA* and *narB)* and dissimilatory (DNRA) reductase genes (*nirBD and narIV)* were differentially abundant in burned plots (p_adj_ < 0.05; Fig. 5c, 7). In contrast, denitrification genes (*nirK*, *narGHYZ*) were differentially abundant in burned plots at early post-fire time points (17 and 25 days; p_adj_< 0.05; Fig. 5c, 7). However, genes associated with N□O and N□ production were not differentially abundant at any time point throughout the year. Additionally, nitrification genes (*hao, pmoA-amoA, pmoB-amoB*) were differentially abundant in burned plots at 131 days post-fire (p_adj_ < 0.05; Fig. 5c, 7). Lastly, while urea cycling pathways were overrepresented in unburned plots at 17 and 25 days, they were differentially abundant in burned plots from 17 to 131 days post-fire (p_adj_ < 0.05; Fig. 5d, 7).

**Figure 7.**
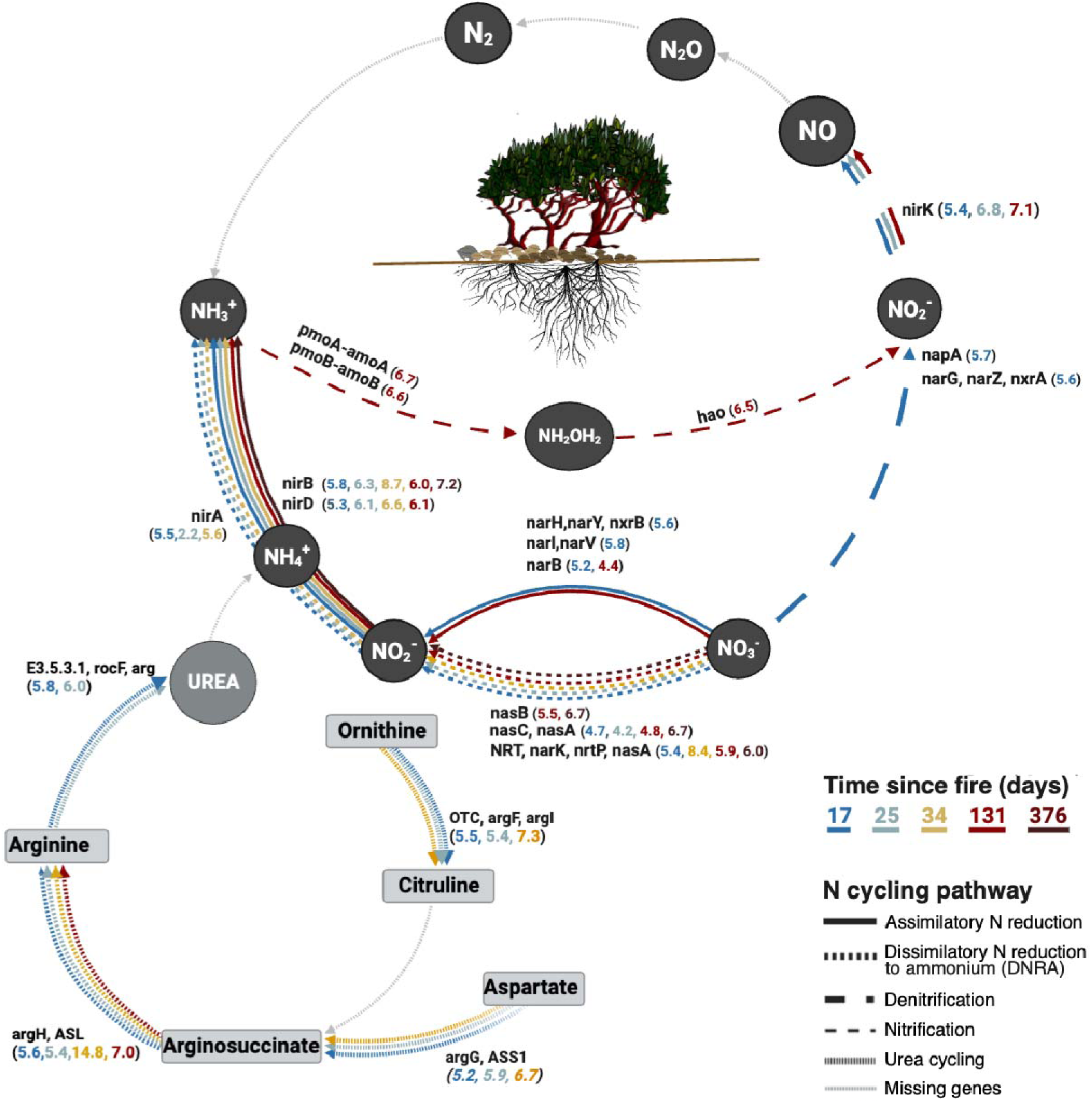
The nitrogen cycle with arrows indicating genes that were differentially represented in the burned treatment at each time point (17, 25, 34, 131, and 376 days post-fire; Wald’s test in DESeq2; *p*<0.05). Line type represents the different nitrogen cycling pathways, colored by timepoint and gray solid lines represent pathways that were not expressed during the first year post-fire. Numbers in parentheses represent the mean log2fold change (DESeq2) for each represented gene and are colored by the timepoint in which that gene was significantly abundantly represented.

#### 1.3.6 PyOM genomic potential of dominant taxa differed in burned versus unburned soils

Genes for catechol and protocatechuate ortho-cleavage and the β-ketoadipate pathway (*fad* genes) were overrepresented in MAGs from burned plots, particularly within Proteobacteria, Actinobacteria, and Acidobacteriota (Fig. 8; Supplementary file 1c). Pyrophilous *Massilia*, which dominated the 16S rRNA gene sequences in burned plots across time (Fig. S5b), was represented by MAGs Holy_71 and Holy_91 (Fig. 8) and encoded genes for catechol ortho-cleavage (*catABC*) and the β-ketoadipate pathways (*pcaDLJ, fadA*). The protocatechuate ortho-cleavage pathway (*pcaCH*) was encoded by *Noviherbaspirillium* (MAGs Holy_91, Holy_361, Holy_395, and Holy_399), which dominated the burned plots at 131 and 376 days (Fig. S4b), the Burkholderiales-SG8-41 Solirubrobacteraceae (MAG Holy_399) and Chthoniobacterales-AV80 (MAG Holy_206; Fig. 8).

**Figure 8.**
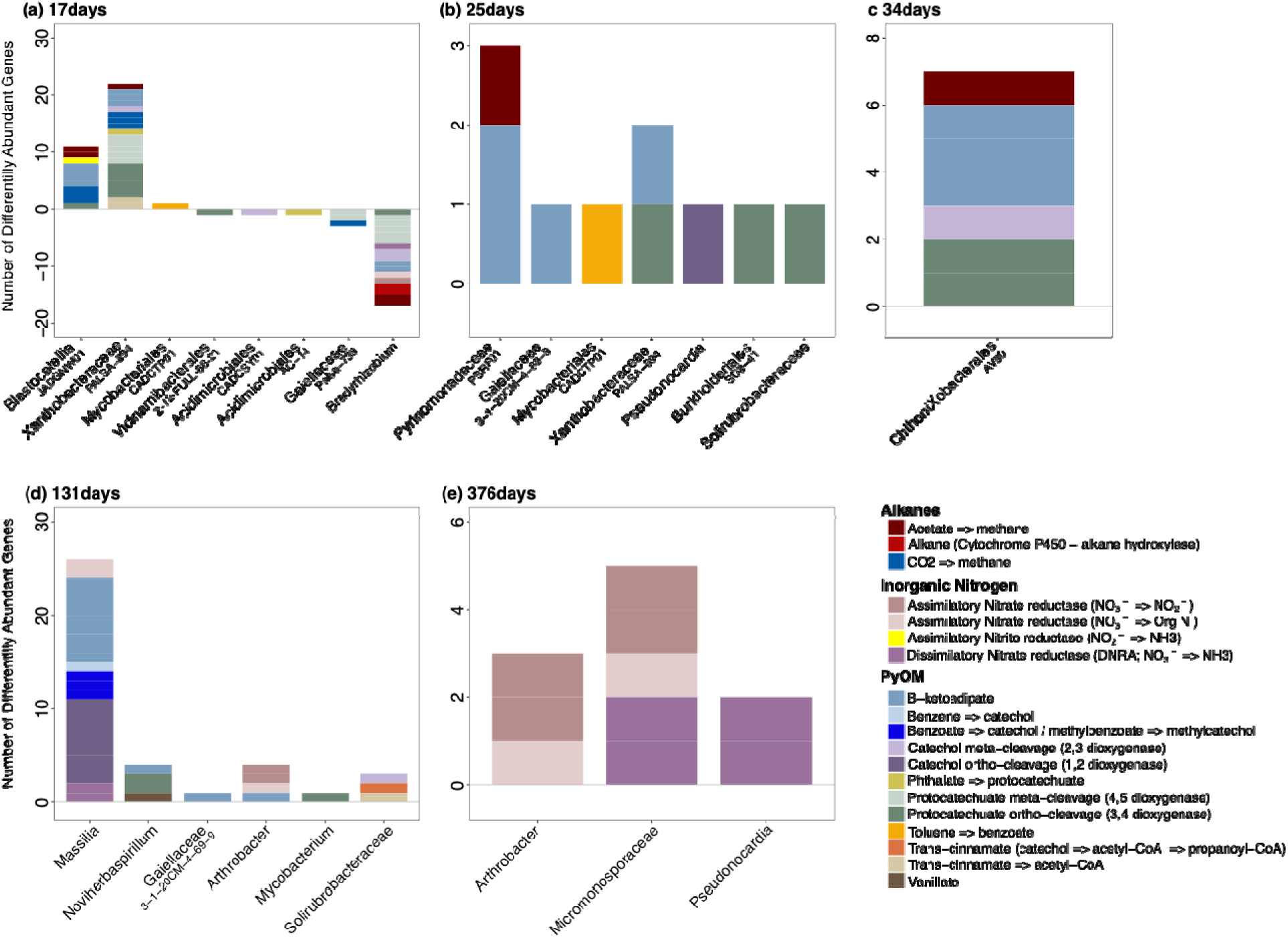
Taxonomic annotation for the most dominant, differentially abundant PyOM, inorganic nitrogen and organic N (urea) genes based on DESeq analysis (adjusted P value < 0.05 and absolute log2 fold-change >0). Taxonomy based on blasting contigs against the Metagenome Assembled Genomes (MAGs) represented in the x-axis.

Catechol meta-cleavage genes (*dmpBC*, *xylEG*) were encoded by MAGs for Proteobacteria, Xanthobacteraceae-PALSA-894 (MAG Holy_368) and Verrucomicrobiota, Chthoniobacterales-AV80 (*catE*, MAG Holy_206; Fig. 8). Lastly, protocatechuate meta-cleavage pathway genes (*ligClK, galC*) in burned plots were differentially abundant by *Noviherbaspirillium* (MAGs Holy_91, 381 and 395) and Xanthobacteraceae*-PALSA-894* (MAG Holy_368; Fig. 8). In contrast, unburned plots were enriched for lignin degrading genes (*pcaD, fadAl, praC, xylH, pracH)*, at 17-25 days, encoded by MAGs representing *Bradyrhizobium* (MAG Holy_369) and Acidimicrobiales-AC-14 (MAG Holy_399).

#### 1.3.7 Nitrogen genomic potential of dominant taxa differed in burned versus unburned soils

Genes encoding NO_3_^-^ retention pathways, including assimilatory NO_3_^-^ reductase (*nasABC, nirA*) and DNRA (*nirBD*), were differentially abundant in MAGs representing pyrophilous taxa within the Actinobacteriota, such as *Arthrobacter* (*nasABC*; MAG Holy_114), Micromonosporaceae (*nasABC*; *nirBD*; MAG Holy_395) and *Pseudonocardia* (*nirBD*; MAG Holy_362; Fig. 8). The Proteobacteria *Massilia* (MAG Holy_71) also encoded genes for N retention, including DNRA (MAG Holy_71; *nirD*) and NO_3_^-^ assimilation (*NRT, narK, nrtP, nasA*; Fig. 8). Additionally, *Arthrobacter* (MAG Holy_114) and the family *Micromonosporaceae* (MAG Holy_395) encoded genes for NO□□ assimilation retention pathways (*NRT, narK, nrtP, nasA*; Fig. 8). In contrast, unburned soils had significant abundances of NO□□ assimilation, NO□□ dissimilatory reductase, and denitrification pathways only at 17 days post-fire (Fig. 8), encoded by non-pyrophilous genera particularly *Bradyrhizobium* (MAG Holy_369, Holy_46; NRT, narK, nrtP, nasAC, napA; Fig. 8).

### 1.4 Discussion

Wildfire significantly altered the genomic functional potential of post-fire soil bacteria across five timepoints, spanning from two weeks to one year post-fire, mirroring significant shifts in bacterial community composition (Pulido-Chavez et al., 2023). Notably, the genomic potential for PyOM degradation increased progressively over time in the burned but not unburned plots, with catechol and protocatechuate ortho-cleavage (i.e., β-ketoadipate) pathways emerging as the primary bacterial mechanisms for PyOM degradation. Concurrently, N cycling genes increased over time in burned plots, with a notable increase in genes for nitrification and a shift towards genes associated with N retention, such as facilitating the conversion of NO□□ to microbial biomass and NO□□ to NH□. Finally, the increase in functional genes for PyOM degradation and N cycling aligned with the dominance of pyrophilous bacterial genera, including *Massilia* and *Noviherbaspirillum*, each encoding distinct pathways for PyOM and N cycling. These findings suggest that resource acquisition traits drive bacterial secondary succession in post-fire soils, underscoring the critical role of pyrophilous microbes in shaping soil C and N dynamics and enhancing post-fire functional restoration.

#### 1.4.1 Bacterial functional capacities shift from labile C to PyOM across succession

We previously hypothesized that post-fire microbial succession transitions from an initial dominance of thermotolerant microbes to fast-colonizers, culminating in microbes capable of exploiting post-fire resources, such as N and PyOM (Pulido-Chavez et al., 2023). Using metagenomes, we corroborated this hypothesis, revealing increased abundance of PyOM degradation genes in burned but not unburned plots over time, coinciding with the recovery of bacterial species richness and diversity. This suggest that successful post-fire genera require metabolic pathways for PyOM degradation to establish and dominate the post-fire system.

Post-fire succession involved trade-offs in C use, with early colonizers favoring labile C and later taxa shifting to more complex, aromatic C sources. During the first 17-25 days, bacteria encoding genes for acetate metabolism, a prevalent post-fire labile C source (Blank et al., 1994; Certini, 2005; Kutscha and Pflugl, 2020) became dominant (*ACSS1_2, acs* and *pta)*, indicating that early successional bacteria are well-equipped to degrade labile C sources to support rapid growth and dominance. As labile C diminished (Fig. 5), bacterial communities transitioned to taxa capable of degrading complex aromatic C sources like PyOM (Fig. 8) (Pulido-Chavez et al., 2023). Indeed, by 131 days post-fire, taxa encoding multiple PyOM degradation genes, such as *Noviherbaspirillum* and *Massilia* became dominant, supporting findings from studies using PyOM treatments (Chen et al., 2021; Cheng et al., 2022; Zeba et al., 2023) and incubation experiments (Song et al., 2017; Woolet and Whitman, 2020) that highlight these genera’s role in using PyOM as a C source.

In our study, *Noviherbaspirillium* increased in relative abundance over time, dominating 16S sequences at 131 and 376 days post-fire, while *Massilia* remained consistently dominant throughout the year (4). The timing difference likely reflects their distinct PyOM degradation strategies. *Massilia* uses the simpler catechol and β-ketoadipate ortho-cleavage pathway, providing immediate access to C resources, which may explain its early and continuous dominance. In contrast, *Noviherbaspirillum* relies on the more complex protocatechuate ortho- and meta-cleavage pathway, which is a slower process due to its regulatory complexities (Johnson and Beckham, 2015). Since the protocatechuate pathway is also associated with a growth lag and delayed substrate utilization (Johnson and Beckham, 2015), this could potentially explain the delayed emergence of *Noviherbaspirillum*, which might take longer to adapt and begin degrading PyOM effectively. These findings suggest that the success of microbial taxa post-fire depends on their ability to use different C sources, with some taxa adapting more quickly to available resources due to their metabolic pathways. Together, these findings suggest that microbial succession and dominance in post-fire environments are influenced by the functional potential for distinct degradation pathways for labile C and PyOM overtime.

Previous research suggest interspecies resource competition between pyrophilous bacteria and fungi, with bacteria encoding the catechol ortho-cleavage pathway and fungi encoding the protocatechuate ortho-cleave pathway (Fuchs et al., 2011). Our bacterial-optimized bioinformatic pipelines yielded no fungal MAGs, but our ASV dataset captured fungal ASVs throughout the year (Pulido-Chavez et al., 2023), highlighting a methodological limitation. Indeed, transcriptome data show that *Aspergillus*, a dominant fungal genus in our ITS2 data (Pulido-Chavez et al., 2023), encodes PyOM degradation genes via the β-ketoadipate pathway (Sgro et al., 2023). Thus, we hypothesize that fungi contribute additional PyOM degradation genes, requiring bioinformatic advances to fully explore this potential.

#### 1.4.2 Post-fire chaparral bacteria preferentially degrade PyOM via the ortho-cleavage degradation pathway

The greater number of differentially abundant PyOM degradation genes in burned compared to unburned plots, together with the dominance of genes for the complete catechol (*catABC*) and protocatechuate (*pcaHGBCDLJ*) ortho-cleavage pathways in burned soils throughout the year, suggests that post-fire bacteria may preferentially use ortho-cleavage over meta-cleavage pathways. This aligns with findings from post-fire coniferous forests, where over 50% of MAGs contained the ortho-cleavage pathway (Nelson et al., 2022), a pathway also enriched following oil spills, which similarly contain high levels of aromatic compounds (Brock Melissa et al., 2025). There results suggest that high-severity fires, which generate substantial PyOM or aromatics (Singh et al., 2012), may favor this pathway in both chaparral and forest biomes. Indeed, the ortho-cleavage pathway is the primary route for aerobic aromatic degradation (Granja-Travez et al., 2020; Weiland et al., 2022), known for its efficiency in processing diverse aromatic compounds (Bugg et al., 2011), converting C into biomass (Basha et al., 2010; Aghapour et al., 2013), and supporting rapid bacterial growth (Song et al., 2002). However, while both catechol and protocatechuate ortho-cleavage pathways are widespread across bacterial species, their gene arrangements vary (Parke et al., 2000; Buchan et al., 2004), which can influence how effectively bacteria break down and utilize PyOM as a C source in different substrates. Here, we found that dominant Proteobacteria MAGs, such as *Noviherbaspirillum* and *Massilia* along with other MAGs including Xanthobacteraceae PALSA-89, and the Acidobacteriota Chthoniobacterales AV80 and Solirubrobacteraceae, contain genes for catechol (*catABC*) and protocatechuate (*pcaHGBCDLJ*) ortho-cleavage degradation pathways. Additionally, we found a timing difference in *Massilia* and *Noviherbaspirillum* emergence (17 days vs 131 days), which supports the idea that their distinct PyOM degradation strategies may reflect variations in gene arrangements within these pathways, influencing their ability to utilize PyOM as a C source. However, further research is needed to explore their specific gene arrangements. Together, our results suggest that pyrophilous bacteria in wildfire-adapted ecosystems, like chaparral, may have evolved a preference for the easier-to-degrade ortho-cleavage pathway to convert PyOM into biomass, thereby facilitating the exploitation of PyOM as a C source and promoting microbial secondary succession

#### 1.4.3 Post-fire microbiomes express genes associated with inorganic N retention and nitrification

We found a notable increase in the genomic potential for nitrification and N retention over time in burned soils. As bioavailable N diminished, bacterial communities likely transitioned to taxa specialized in N retention, potentially favoring increases in biomass, which is consistent with the dominance of *Massilia* MAGs, which accounted for ∼ 60% of 16S sequences throughout the year (Pulido-Chavez et al., 2023) and encoded key genes for N retention (Assimilatory: *nasABC, NRT, narK, nrtP*, *nirA* and *narB* and DNRA: *nirBD and narIV*). By 131 days post-fire, succession was marked by the emergence of bacterial genera *Arthrobacter*, *Blastocatellia*, and Micromonosporaceae MAGs, which exhibited high genomic potential for N immobilization via the assimilatory NO□□ reduction pathway. These findings support observations of microbial N immobilization in post-fire environments (Xu et al., 2022), including chaparral (Goodridge et al., 2018) and studies showing the enrichment of key N retention genes (*nirB/D*) up to four years post-fire (Dove et al., 2022). Moreover, studies involving PyOM additions have shown that PyOM facilitates N retention in burned soils (Bai et al., 2015; Yu et al., 2023), suggesting that post-fire N retention by altering microbiomes may represent a common strategy for stabilizing nutrient cycling in fire disturbed ecosystems. Additionally, *Massilia, Pseudonocardia*, and Micromonosporaceae have the capacity to reduce NO□□ to NH□□ via dissimilatory NO□□ reductase (*nirB/D*), trading off a mobile and reactive anion for a cation more often retained on soil exchange sites (Kalisz and Stone, 1980; Homyak et al., 2021). The observed enrichment of N retention genes over time and the presence of microbial taxa capable of N immobilization and assimilation, align with the known N reducing and assimilating functions of *Massilia* and *Arthrobacter* (Cacciari et al., 1986; Bailey et al., 2014; Liu et al., 2023b).

#### 1.4.4. Denitrification and urea genes increase in early post-fire chaparral

Nitrogen losses from volatilization during fires and leaching afterward often surpass the ecosystem’s capacity to sequester N (Turner et al., 2007; Stephens and Homyak, 2023), especially if fires disrupt N-cycling microbes (Knicker, 2007; Hanan et al., 2016). Our findings indicate that the post-fire surge in bioavailable N (Wan et al., 2001) stimulates bacterial activity, leading to an overrepresentation of denitrification genes (*nirK, narG, narZ*) during early secondary succession (17-25 days). However, genes for complete nitrate reduction to N_2_ via denitrification (NO → N□O → N□) were notably rare, despite studies reporting production of N_2_O and N_2_ in post-fire environments (Stephens and Homyak, 2023). While we could not detect the full suite of denitrification genes, complete reduction to N_2_ and partial reduction to N_2_O may have still occurred, but may have gone undetected because we mostly sampled during dry conditions that constrain denitrification (Krichels et al., 2023). Nevertheless, our measurements suggest that early post-fire bacteria contain a high number of denitrification genes, potentially creating conditions that accelerate N loss via gaseous pathways. Additionally, urea cycling pathways were enriched in burned plots from 17-to-131 days post-fire, but only from 17-to-25 days in unburned plots. This suggest that post-fire microbes exploit urea as a N source, converting it to NH_4_^+^ for growth, thus favoring microbes with urea cycling genes in the N rich environment. Together, our results show that bacteria with specialized N metabolism dominate and mediate post-fire succession, transitioning from denitrification to nitrification and ultimately expressing genes for N retention as bioavailable N declines.

### 1.5 Conclusions

Our high-resolution gene-centric metagenomic analysis reveals rapid enrichment of genomic functional potential for PyOM, alkanes, and inorganic N in burned soils over the first post-fire year. These functional shifts align with the dominance of pyrophilous bacteria, such as *Massilia* and *Noviherbaspirillium,* which encode pathways for ortho-cleavage catechol and protocatechuate degradation, while potentially assimilating N into biomass. This suggest that resource acquisition traits are key drivers of bacterial secondary succession in fire-affected soils. The incomplete genomic potential for PyOM and N pathways within a single bacterial genus, combined with distinct degradation routes employed by different bacteria, suggests metabolic handoffs between groups. Specifically, some taxa degrade complex compounds, while others utilize byproducts, indicating widespread resource transfers in wildfire-affected communities. While the ability of post-fire bacteria to capitalize on PyOM and N has been previously hypothesized (Woolet and Whitman, 2020; Pulido-Chavez et al., 2023), and shown for some taxa (Cobo-Díaz et al., 2015; Fischer et al., 2021; Nelson et al., 2022), our research provides the first direct evidence supporting our conceptual model of trait trade-offs among bacteria (Enright et al., 2022), demonstrating that specific functional traits drive bacterial succession in fire-impacted soils. These findings have significant implications for predicting how bacterial communities influence C sequestration and N cycling in post-fire ecosystems. Moreover, by revealing previously unrecognized functional shifts in bacterial communities across a high-resolution successional timeline, we fill a critical gap in post-fire ecological research and provide essential insights into the long-terms impact of fire on microbial-mediated ecosystem processes.

## 1.6 Acknowledgments

We thank the Joint Genome Institute (JGI) Community Sequencing Project (CSP) New Investigator Award 507364 to SIG, the Bureau of Land Management (BLM) Joint Fire Science Program (JFSP) Graduate Research Innovation (GRIN) Award 012641 002 to MFPC and SIG, the United States Department of Agriculture (USDA)-NIFA Award #2022-67014-36675 to SIG and PMH, and the Department of Energy (DOE) BER Award #DE-SC0023127 to SIG, PMH and MJW for funding this project. We further thank the Cleveland National Forest and the Trabuco Ranger District, including District Ranger Darrel Vance and Emily Fudge, Jeffrey Heys, Lauren Quon, Jacob Rodriguez, and Victoria Stempniewicz, for their help with permitting. We thank JGI Metagenome Program, including Nicole Shapiro, Natasha Brown, Emiley Eloe-Fadrosh, and Simon Roux, for their help in processing and submitting our metagenomic samples. Lastly, we thank Judy A. Chung, Aral C. Greene, Elizah Stephens, Dylan Enright, and Sameer S. Saroa for fieldwork assistance and James Randolph for molecular assistance. We thank Jason Stajich and anonymous reviewers for helpful feedback on the manuscript.

## 1.8 Data Availability Statement

The metagenomic reads and bacterial MAGs have been deposited to the National Center for Biotechnology Information Sequence Read Archive under BioProject PRJNA761539. Amplicon sequence reads for bacterial 16S rRNA and fungal ITS2 have been deposited to the National Center for Biotechnology Information Sequence Read Archive under BioProject PRJNA761539. All statistical R scripts/codes are available on GitHub and include the sample metadata required for analysis https://github.com/pulidofabs/Chaparral-Metagenomes.

