## SupplementaryData for "Chaparral wildfire shifts the functional potential for soil pyrogenic organic matter and nitrogen cycling"

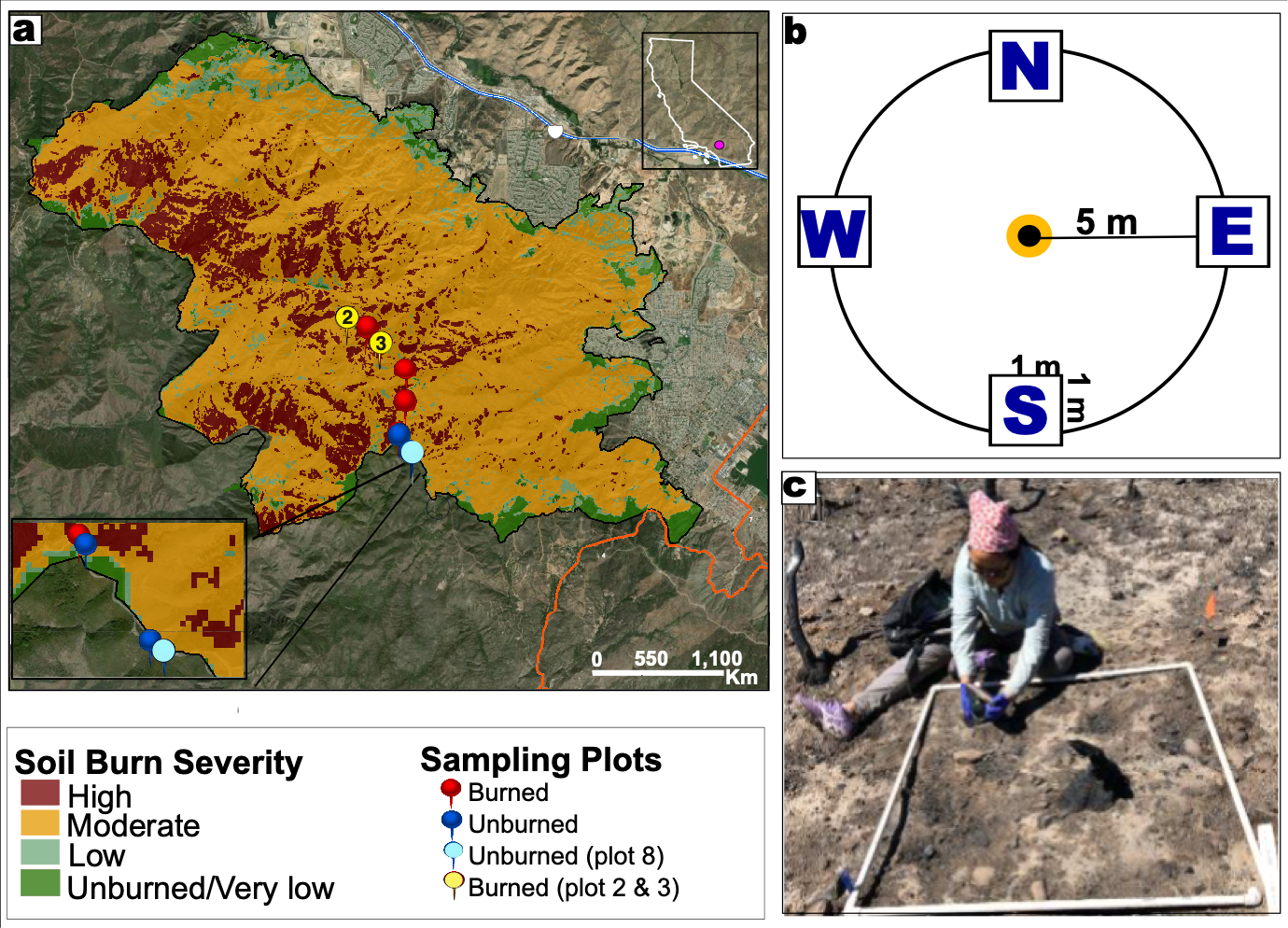

**Figure S1.** A) Burned Area Emergency Response (BAER) map of the soil burn severity within the 2018 Holy Fire including our 6 burned and 3 unburned plots. B) sampling design for plots with four 1m^2^ subplots at 5m from the center in each cardinal direction. C) Image of collection top 10 cm of soil beneath the ash in a 1-m^2^ subplot immediately after fire. For metagenomic analysis, burned plots 2 and 3 (yellow pins) and unburned plot 8 (light blue pin) were used. Figure adapted from (Pulido-Chavez et al., 2023)**.**

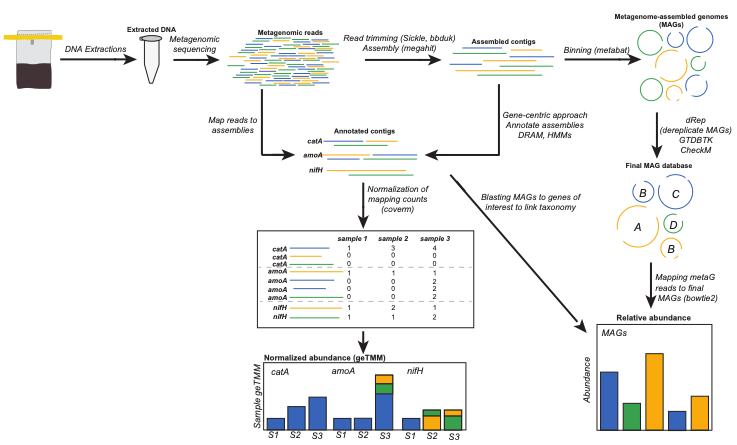

**Figure S2.** Methodological map for the bioinformatic steps used to analyze Illumina NovaSeq shotgun metagenomic sequencing.

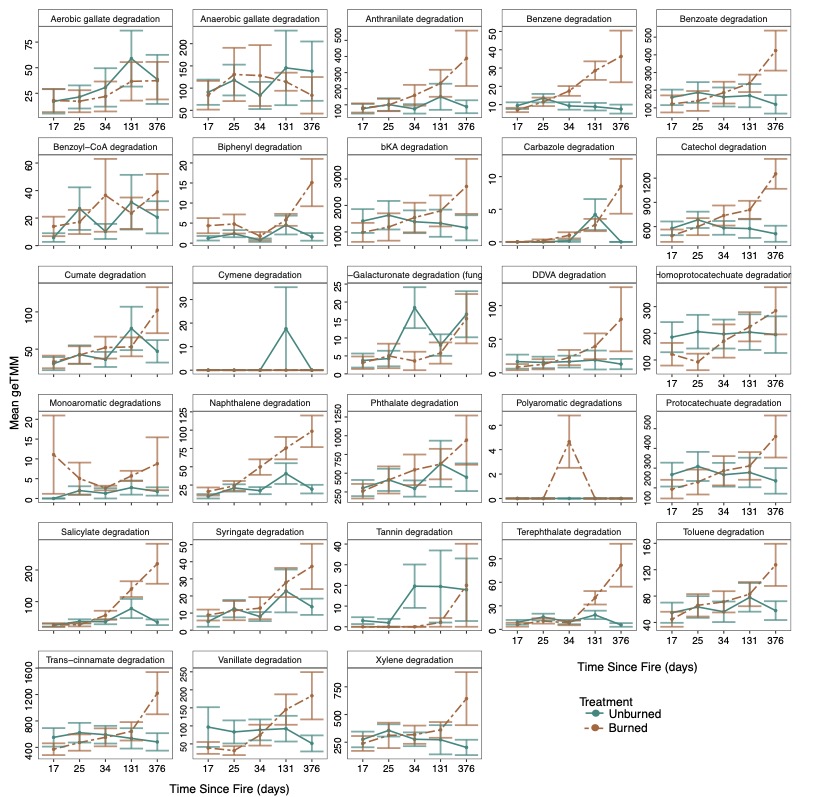

**Figure S3.** Effect of time (days since fire) on Pyrogenic Organic Matter (PyOM) degradation pathways in burned (brown) and unburned (blue-green) plots. The mean geTMM values represent contigs normalized by contig length, adjusted using the trimmed mean of M-values (TMM), and further normalized by library depth and gene length.

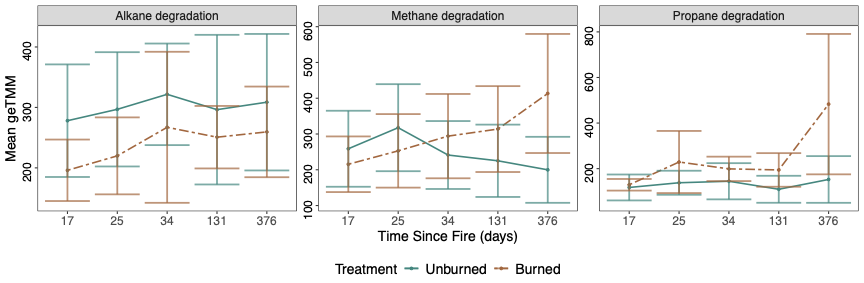

**Figure S4.** Effect of time (days since fire) on alkane degradation pathways (readily available C sources) in burned (brown) and unburned (blue-green) plots. The mean geTMM values represent contigs normalized by contig length, adjusted using the trimmed mean of M-values (TMM), and further normalized by library depth and gene length.

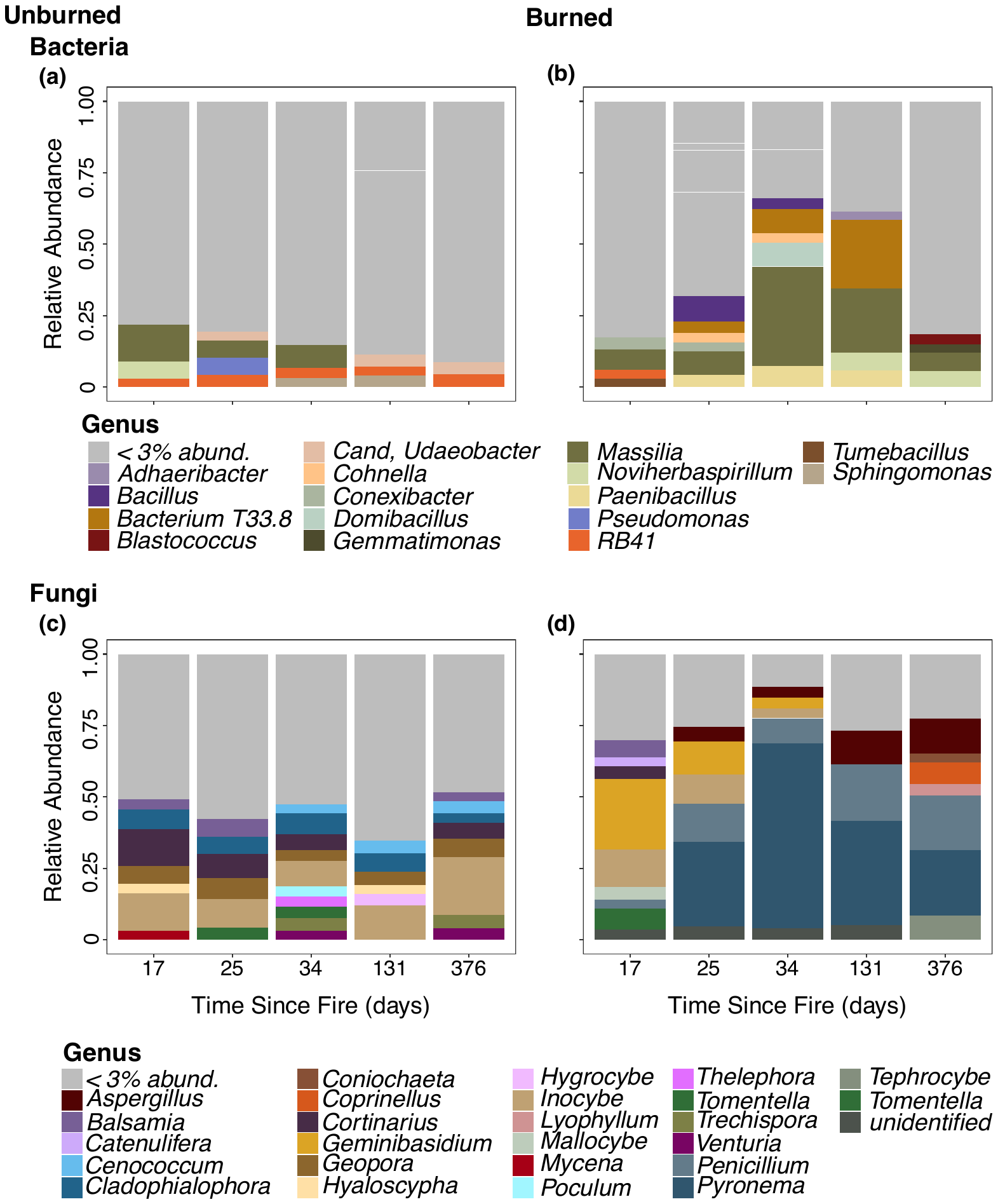

**Figure S5.** Relative abundance of bacterial and fungal communities across five timepoints (17, 34, 67, 131, and 376 days post-fire). For a detailed description of relative abundance trends over the entire year (9 timepoints), refer to Pulido-Chavez et al. (2023).

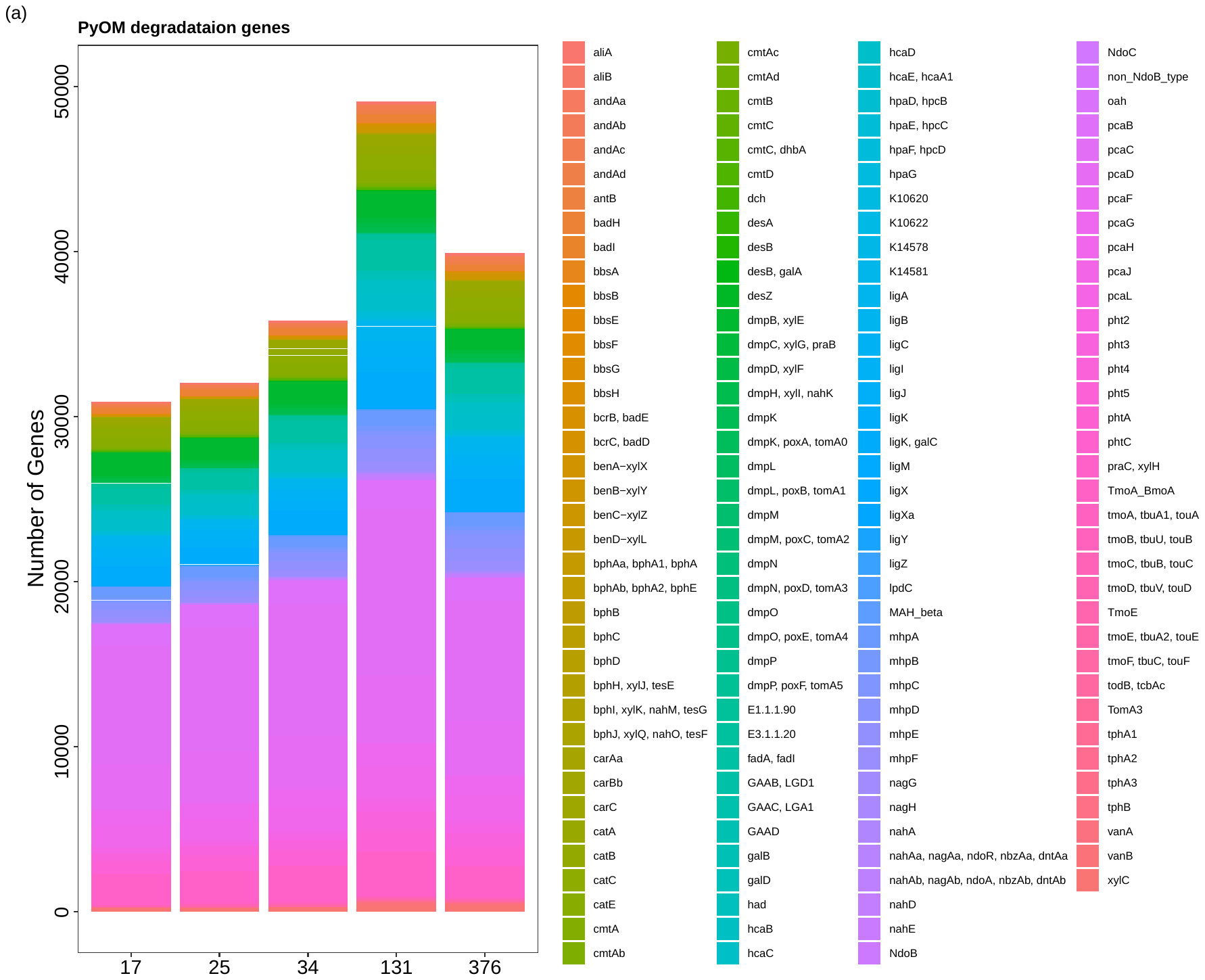

**Figure S6.** Total number of genes for Pyrogenic Organic Matter (PyOM) degradation across time (Time since fire in days on the x-axis) in the burned plots for the entire dataset.

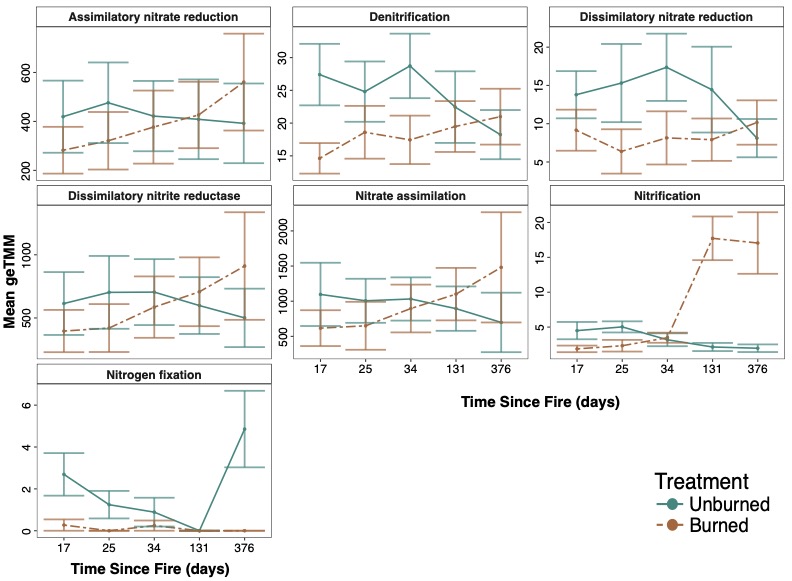

**Figure S7.** Effect of time (days since fire) on inorganic nitrogen cycling pathways in burned (brown) and unburned (blue-green) plots. The mean geTMM values represent contigs normalized by contig length, adjusted using the trimmed mean of M-values (TMM), and further normalized by library depth and gene length.

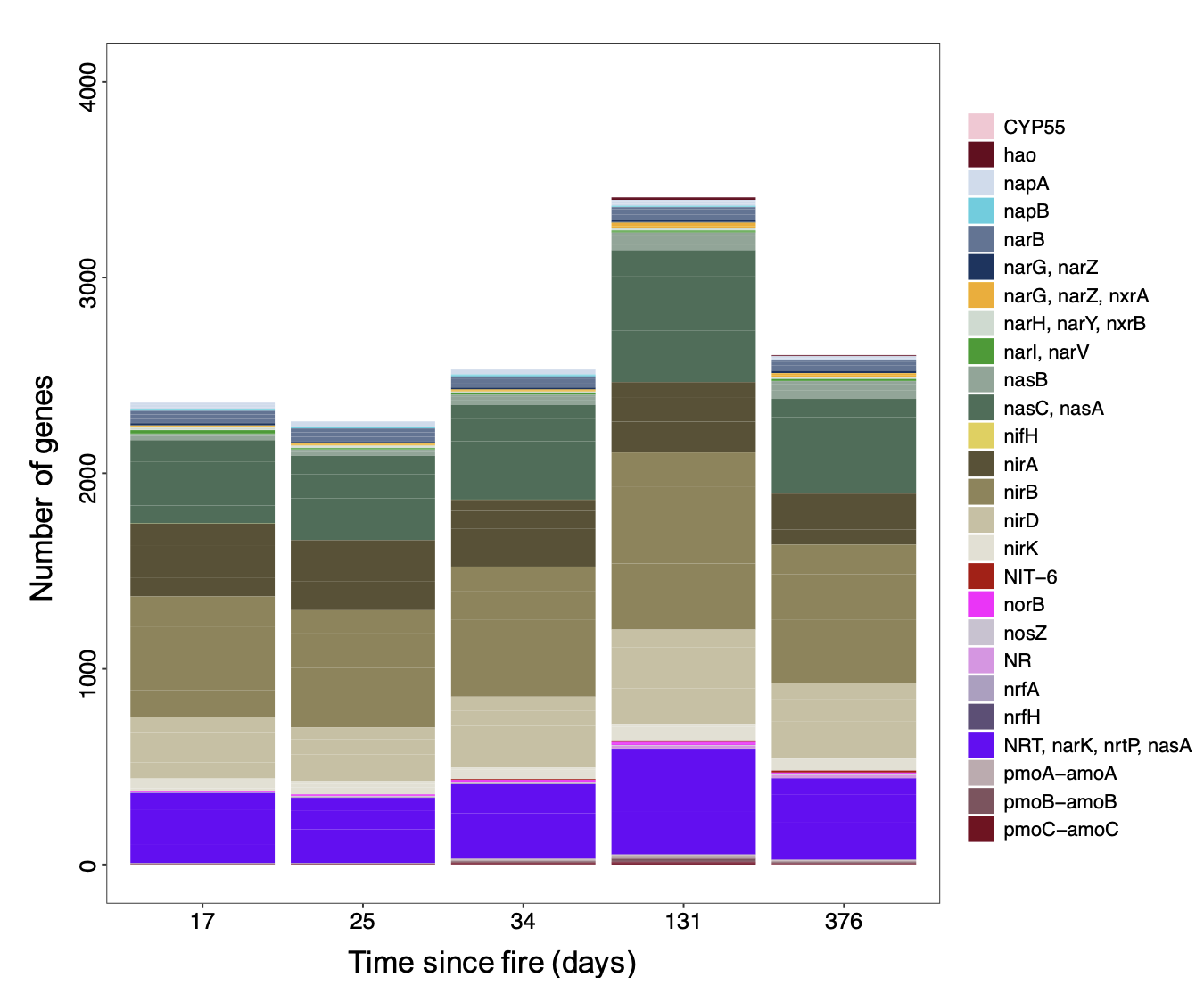

**Figure S8.** Total number of genes (legend) for inorganic nitrogen cycling across time (time since fire in days) in the burned plots.

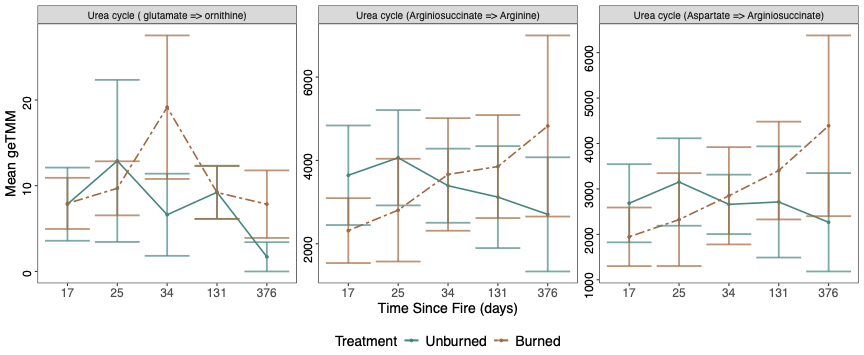

**Figure S9** Effect of time (days since fire) on organic nitrogen cycling pathways in burned (brown) and unburned (blue-green) plots. The mean geTMM values represent contigs normalized by contig length, adjusted using the trimmed mean of M-values (TMM), and further normalized by library depth and gene length.

**Table S1.** Permanova results of the effects of treatment (Fire) and Time (Time since fire in days) on alkane, PyOM, inorganic nitrogen, and organic nitrogen (urea) gene composition. Significance based on p<0.05 and based on Bray-Curtis dissimilarity.

| **Functional Gene** | **Variable** | **Sum of Sqs** | **R2** | **F** | **Pr (>F)** |
| --- | --- | --- | --- | --- | --- |
| **alkanes** | Fire | 1.47 | 0.16 | 5.59 | **0.001** |
|  | Time | 0.51 | 0.06 | 1.95 | **0.02** |
|  | Fire x Time | 0.43 | 0.05 | 1.63 | 0.07 |
| **PyOM** | Fire | 1.39 | 0.15 | 5.25 | **0.001** |
|  | Time | 0.49 | 0.05 | 1.84 | **0.03** |
|  | Fire x Time | 0.47 | 0.05 | 1.76 | **0.03** |
| **inorganic nitrogen** | Fire | 1.51 | 0.16 | 5.65 | **0.001** |
|  | Time | 0.45 | 0.05 | 1.69 | **0.04** |
|  | Fire x Time | 0.53 | 0.06 | 1.97 | **0.02** |
| **Urea (organic nitrogen)** | Fire | 1.39 | 0.15 | 5.35 | **0.001** |
|  | Time | 0.47 | 0.05 | 1.83 | **0.02** |
|  | Fire x Time | 0.47 | 0.05 | 1.81 | **0.04** |

**Table S2.** Permutest results for the betadisper analysis for inorganic nitrogen, organic nitrogen (urea), PyOM and alkane pathways, based on 9999 permutations and “plot” as random effect.

|  |  |  | **Sum Sq** | **Mean Sq** | **F** | **Pr(>F)** |
| --- | --- | --- | --- | --- | --- | --- |
| **Inorganic** | Burn | Groups | 0.07 | 0.02 | **3.82** | **0.04** |
| **nitrogen** |  | Residuals | 0.04 | 0.004 |  |  |
|  | Unburned | Groups | 0.01 | 0.003 | 0.52 | 0.7 |
|  |  | Residuals | 0.06 | 0.01 |  |  |
| **urea (organic nitrogen)** | Burn | Groups | 0.08 | 0.02 | **4.51** | **0.03** |
|  |  | Residuals | 0.04 | 0.004 |  |  |
|  | Unburned | Groups | 0.01 | 0.003 | 0.52 | 0.7 |
|  |  | Residuals | 0.06 | 0.01 |  |  |
| **PyOM** | Burn | Groups | 0.1 | 0.02 | **5.32** | **0.02** |
|  |  | Residuals | 0.05 | 0.005 |  |  |
|  | Unburned | Groups | 0.01 | 0.002 | 0.39 | 0.8 |
|  |  | Residuals | 0.06 | 0.01 |  |  |
| **alkane** | Burn | Groups | 0.08 | 0.02 | **4.65** | **0.02** |
|  |  | Residuals | 0.04 | 0.004 |  |  |
|  | Unburned | Groups | 0.01 | 0.003 | 0.5 | 0.7 |
|  |  | Residuals | 0.05 | 0.01 |  |  |
| DF (4), Residuals (10), N permutations (9999) per pathway | | | | | | |

**Table S3.** Generalized negative binomial results showing the effects of fire, time since fire (in days), and their interaction on nitrogen, urea, pyrogenic organic matter (PyOM), and alkane cycling genes. Significant effects (p < 0.05) are highlighted in bold. Negative estimate and z-value scores indicate that time leads to a decrease in the inorganic nitrogen cycling pathways.

| **Nitrogen** | | | | **Urea** | | |
| --- | --- | --- | --- | --- | --- | --- |
|  | **Estimate** | **Std. Error** | **Pr(>\|z\|)** | **Estimate** | **Std. Error** | **Pr(>\|z\|)** |
| (Intercept) | 4.57 | 0.26 | **<2e-16** | 7.09 | 0.26 | **<2e-16** |
| Fire | 0.02 | 0.32 | 0.96 | 0.11 | 0.31 | 0.71 |
| Time | -0.19 | 0.1 | 0.06 | -0.16 | 0.07 | **0.02** |
| Fire x Time | 0.37 | 0.16 | **0.02** | 0.28 | 0.01 | **<2e-16** |
| **Random Effects** |  |  |  |  | |  |
| Time/Subplot |  | | |  | | |
| Marginal/Conditional R2 | 0.014/0.11 | | | 0.08/ 0.99 | | |
| **PyOM** |  |  |  | **Alkane** | |  |
|  | **Estimate** | **St. Error** | **Pr(>\|z\|)** | **Estimate** | **St. Error** | **Pr(>\|z\|)** |
| (Intercept) | 5.63 | 0.29 | **< 2e-16** | 5.63 | 0.29 | **< 2e-16** |
| Fire | 0.28 | 0.34 | 0.41 | 0.28 | 0.34 | 0.41 |
| Time | -0.19 | 0.07 | **3.00E-03** | -0.19 | 0.07 | **0.003** |
| Fire x Time | 0.38 | 0.07 | **4.20E-08** | 0.38 | 0.07 | **4.20E-08** |
| **Random Effects** |  |  |  |  | |  |
| Time/Subplot |  | | | 0.013/0.31 | | |
| Marginal/Conditional R2 | 0.026/0.18 | | | 0.03/ 0.18 | | |

**Table S4.** Generalized negative binomial regression for the effects of each individual time point (Time since fire in days) on nitrogen and urea cycling genes (top panel) and PyOM and alkane genes (bottom panel) for burned plots. Significance based on p<0.05, denoted in bold. Negative estimate and z-value scores indicate that time leads to a decrease in the inorganic nitrogen cycling pathways.

| **Inorganic nitrogen** | | | |  | **Urea (organic nitrogen)** | | |
| --- | --- | --- | --- | --- | --- | --- | --- |
|  | **Est** | **Std. Error** | **p** |  | **Est** | **Std.**  **Error** | **p** |
| (Intercept) | 4.64 | 0.34 | **<2e-16** |  | 7.19 | 0.25 | **<2e-16** |
| Fire | -0.35 | 0.46 | 0.45 |  | -0.52 | 0.31 | 0.09 |
| Time 25d | 0.31 | 0.35 | 0.37 |  | 0.23 | 0.01 | **<2e-16** |
| Time 34d | -0.08 | 0.33 | 0.81 |  | -0.18 | 0.01 | **<2e-16** |
| Time 131d | -0.24 | 0.33 | 0.46 |  | -0.21 | 0.01 | **<2e-16** |
| Time 376d | -0.42 | 0.34 | 0.21 |  | -0.38 | 0.01 | **<2e-16** |
| Fire : Time 25d | -0.16 | 0.52 | 0.76 |  | 0.39 | 0.03 | **<2e-16** |
| Fire : Time 34d | 0.35 | 0.53 | 0.51 |  | 0.69 | 0.02 | **<2e-16** |
| Fire : Time 131d | 0.70 | 0.48 | 0.14 |  | 0.92 | 0.02 | **<2e-16** |
| Fire : Time 376d | 0.99 | 0.50 | **0.05** |  | 1.04 | 0.02 | **<2e-16** |
| **Random Effects** | | | | | | | |
| Plot | 0.24 | | |  |  | | |
| Marginal/conditional R2 var | 0.021/ 0.12 | | |  |  | | |
| **PyOM** | | | |  | **alkane** | | |
|  | **Est** | **Std. Error** | **p** |  | **Est** | **Std. Error** | **p** |
| (Intercept) | 5.76 | 0.30 | **< 2e-16** |  | 5.76 | 0.30 | **< 2e-16** |
| Fire | -0.28 | 0.37 | 0.45 |  | -0.28 | 0.37 | 0.45 |
| Time 25d | 0.29 | 0.14 | **0.03** |  | 0.29 | 0.14 | **0.03** |
| Time 34d | -0.25 | 0.13 | **6.1E-02** |  | -0.25 | 0.13 | 0.06 |
| Time 131d | -0.33 | 0.13 | **0.01** |  | -0.33 | 0.13 | **0.01** |
| Time 376d | -0.50 | 0.14 | **0.0002** |  | -0.50 | 0.14 | **0.0002** |
| Fire : Time 25d | 0.01 | 0.23 | **9.6E-01** |  | 0.01 | 0.23 | 0.96 |
| Fire : Time 34d | 0.73 | 0.26 | **0.004** |  | 0.73 | 0.26 | **0.004** |
| Fire : Time 131d | 0.78 | 0.19 | **0.0001** |  | 0.78 | 0.19 | **5.3E-05** |
| Fire : Time 376d | 1.25 | 0.22 | **0.000** |  | 1.25 | 0.22 | **2.6E-08** |
| **Random Effects** | | | | | | | |
| Subplot | 0.33 | | |  | Time/Subplot | |  |
| Marginal/conditional R2 var | 0.04 / 0.20 | | |  | 0.04/ 0.20 | | |

**Table S5.** Generalized negative binomial regression for the effects of (time since fire in days) on all the different PyOM cycling (C) pathways for burned and unburned plots independently. Significance based on p<0.05, denoted in bold. Negative estimate and z-value scores indicate that time leads to a decrease in the inorganic nitrogen cycling pathways.

|  | **Burn** | | | | **Unburned** | | | |
| --- | --- | --- | --- | --- | --- | --- | --- | --- |
| **PyOM pathways** |  | **Est.** | **z value** | **P** | **Est.** | **z value** | **P** | **R2 (B),(Un)** |
| **Catechol Meta** | (Int) | 6.29 | 23.54 | **< 2e-16** | 5.99 | 18.27 | **<2e-16** | (0.04/0.21), (0.025/0.31) |
|  | Time | 0.21 | 2.92 | **0.004** | -0.19 | -2.44 | **0.01** |  |
| **Catechol Ortho** | (Int) | 6.56 | 25.14 | **< 2e-16** | 6.26 | 25.08 | **<2e-16** | (0.1/0.29), (0.054/0.28) |
|  | Time | 0.3 | 3.23 | **0.001** | -0.22 | -2.42 | **0.0155** |  |
| **Aerobic Gallate** | (Int) | 2.8 | 8.04 | **8.90E-16** | 3.4 | 8.4 | **<2e-16** | (0.07/0.9), (0.04/0.01) |
|  | Time | 0.17 | 5.06 | **4.30E-07** | 0.24 | 0.73 | 0.463 |  |
| **Anaerobic Gallate** | (Int) | 4.5 | 16.93 | **<2e-16** | 4.49 | 16.64 | **<2e-16** | (0.04/0.3), (0.01/0.61) |
|  | Time | -0.13 | -0.86 | 0.393 | 0.06 | 0.52 | 0.605 |  |
| **Benzene** | (Int) | 2.79 | 10.74 | **< 2e-16** | 1.92 | 5 | **5.70E-07** | (0.13/0.27), (0.05/0.38) |
|  | Time | 0.39 | 3.91 | **9.30E-05** | -0.29 | -2.78 | **0.01** |  |
| **Benzoate** | (Int) | 5.12 | 19.6 | **< 2e-16** | 4.8 | 17 | **<2e-16** | (0.13/0.29), (0.06/0.3) |
|  | Time | 0.38 | 3.72 | **0.0002** | -0.25 | -2.36 | **0.02** |  |
| **BenzoykCoA** | (Int) |  |  |  | 2.92 | 8.46 | **<2e-16** | (0.012/0.01) |
|  | Time |  |  |  | 0.17 | 0.43 | 0.668 |  |
| **Biphenyl** | (Int) | 1.37 | 2.83 | **0.005** | -0.06 | -0.11 | 0.912 | (0.12/0.20), (0.004/0.31) |
|  | Time | 0.53 | 2.02 | **0.04** | -0.11 | -0.34 | 0.734 |  |
| **bka** | (Int) | 7.04 | 25.75 | **<2e-16** | 6.91 | 26.3 | **<2e-16** | (0.06/0.17) |
|  | Time | 0.3 | 2.03 | **0.04** | -0.18 | -1.3 | 0.193 |  |
| **Carbazole** | (Int) | -0.05 | -0.14 | 0.89 | -2.34 | -1.23 | 0.221 | (0.6/0.8), (0.0004/0.96) |
|  | Time | 1.05 | 11.6 | **<2e-16** | 0.06 | 0.4 | 0.691 |  |
| **Cumate** | (Int) | 3.78 | 14.76 | **<2e-16** | 3.47 | 9.45 | **<2e-16** | (0.06/0.13), (0.0003,0.19) |
|  | Time | 0.33 | 2.47 | **0.01** | -0.03 | -0.15 | 0.883 |  |
| **Dgalac** | (Int) | 1.09 | 2.36 | **0.02** | 0.77 | 0.61 | 0.542 | (0.21/0.80), |
|  | Time | 0.46 | 10.14 | **<2e-16** | 0.03 | 0.14 | 0.891 | (1.3e-04/0.82) |
| **DDVA** | (Int) | 3.22 | 11.84 | **<2e-16** | 2.77 | 7.35 | **2.00E-13** | (0.22/0.22), (2.9e-03/2.9e-03) |
|  | Time | 0.7 | 2.35 | **0.02** | -0.08 | -0.21 | 0.837 |  |
| **PCA Meta** | (Int) | 5.87 | 17.62 | **< 2e-16** | 5.96 | 17.32 | **< 2e-16** | (0.11/0.4), (0.03/0.41) |
|  | Time | 0.36 | 5.11 | **3.30E-07** | -0.18 | -2.68 | **0.01** |  |
| **PCA Ortho** | (Int) | 7.89 | 32 | **<2e-16** | 7.75 | 27.51 | **<2e-16** | (0.034/0.22), (0.04/0.33) |
|  | Time | 0.17 | 1.89 | **0.06** | -0.18 | -2.06 | **0.04** |  |
| **Naphthalene** | (Int) | 3.75 | 14.96 | **< 2e-16** | 2.65 | 7.03 | **2.10E-12** | (0.23/0.34), (3.6e-04/0.3) |
|  | Time | 0.56 | 4.67 | **3.00E-06** | -0.02 | -0.16 | 0.871 |  |
| **Phathalete** | (Int) | 6.09 | 32.45 | **<2e-16** | 5.87 | 19.32 | **<2e-16** | (0.43/0.99), (0.003/0.07) |
|  | Time | 0.28 | 73.67 | **<2e-16** | 0.08 | 0.34 | 0.736 |  |
| **Syringate** | (Int) | 2.4 | 5.82 | **5.80E-09** | 1.14 | 1.09 | 0.276 | (0.2/0.5), (0.005/0.92) |
|  | Time | 0.52 | 2.92 | **0.003** | 0.15 | 0.92 | 0.356 |  |
| **Terephthalate** | (Int) | 2.67 | 9 | **<2e-16** | 2.4 | 13.01 | **<2e-16** | (0.6/0.94), (0.05/0.05) |
|  | Time | 0.71 | 31.99 | **<2e-16** | -0.25 | -1.33 | 0.182 |  |
| **Toluene** | (Int) | 4.1 | 15 | **<2e-16** | 3.73 | 10.61 | **<2e-16** | (0.04/0.15) |
|  | Time | 0.26 | 2.3 | **0.02** | -0.14 | -1.2 | 0.232 |  |
| **Trans** | (Int) | 6.18 | 22.61 | **< 2e-16** | 6.07 | 21.76 | **<2e-16** | (0.08/0.20), (0.02/0.2) |
|  | Time | 0.35 | 3.01 | **0.003** | -0.17 | -1.36 | 0.174 |  |
| **Vanillate** | Int) | 4.1 | 13.44 | **<2e-16** | 4.28 | 11.68 | **<2e-16** | (0.36/0.97), (0.04/0.1) |
|  | Time | 0.41 | 34.07 | **<2e-16** | -0.24 | -1.02 | 0.308 |  |
| **Xylene** | (Int) | 5.62 | 19.6 | **< 2e-16** | 5.43 | 24.82 | **<2e-16** | (0.13/0.53), (0.13/0.5) |
|  | Time | 0.26 | 2.62 | **0.01** | -0.24 | -2.53 | **0.01** |  |

**Table S6.** Generalized negative binomial regression for the effects of (time since fire in days) on all the different alkane degradation pathways for burned plots. Significance based on p<0.05, denoted in bold. Negative estimate and z-value scores indicate that time leads to a decrease in the inorganic nitrogen cycling pathways.

|  |  | **Burn** | | | **Unburned** | | |  |
| --- | --- | --- | --- | --- | --- | --- | --- | --- |
| **Alkane pathways** | | **Est** | **z value** | **P** | **Estimate** | **z value** | **P** | **R2**  **(B),(Un)** |
| **methane** | (Int) | 5.51 | 21.43 | <2e-16 | 5.37 | 23.46 | <2e-16 | (0.013/0.07), (0.02/0.07) |
|  | Time | 0.17 | 1.49 | 0.136 | -0.19 | -1.56 | 0.119 |  |
| **alkane** | (Int) | 5.08 | 14.81 | <2e-16 | 5.15 | 10.27 | <2e-16 | (0.0001/0.5), (0.01/0.70) |
|  | Time | -0.01 | -0.1 | 0.919 | -0.12 | -1.15 | 0.252 |  |
| **propane** | (Int) | 5.3 | 19.61 | <2e-16 | 4.47 | 11.76 | <2e-16 | (0.23/0.43), (0.01/0.7) |
|  | Time | 0.37 | 2.21 | **0.0275** | -0.08 | -0.59 | 0.554 |  |

**Table S7.** Generalized negative binomial regression for the effects of time (time since fire in days) on all the different inorganic nitrogen cycling pathways for burned and unburned plots independently. Significance based on p<0.05, denoted in bold. Negative estimate and z-value scores indicate that time leads to a decrease in the inorganic nitrogen cycling pathways.

|  | **Burn** | | | | **Unburned** | | | |
| --- | --- | --- | --- | --- | --- | --- | --- | --- |
|  |  | **Est** | **z value** | **P value** | **Est** | **z value** | **P value** | **R2 (B),(Un)** |
| **Dissimilatory Nitrite** | (Int) | 6.03 | 19.04 | **<2e-16** | 6.15 | 19.21 | < 2e-16 | (0.1/0.99), (0.07/ 0.99) |
|  | Time | 0.2 | 2.06 | **0.0397** | -0.18 | -3.15 | **0.002** |  |
| **Assimilatory Nitrite Reductase** | (Int) | -6.55 | -4.99 | **5.90E-07** | 2.52 | 3.28 | **0.001** | (0.04/1), (0.06/0.73) |
|  | Time | 1.71 | 8.5 | **< 2e-16** | -0.41 | -1.34 | 0.18 |  |
| **Assimilatory Nitrate Reductase** | (Int) | 1.29 | 2.17 | **0.03** | 6.14 | 28.19 | **<2e-16** | (0.0006/0.42), (0.003/0.04) |
|  | Time | -0.03 | -0.19 | 0.85 | -0.07 | -0.35 | 0.72 |  |
| **Dissimilatory Nitrate Reductase** | (Int) | 5.97 | 19.49 | **<2e-16** | 1.99 | 3.55 | **0.0004** | (0.023/0.1), (0.1/0.63) |
|  | Time | 0.18 | 0.93 | 0.35 | -0.44 | -3.24 | **0.001** |  |
| **N assimilation** | (Int) | 6.58 | 22.12 | **<2e-16** | 6.6 | 21.95 | **<2e-16** | (0.15/0.59), (0.14/0.62) |
|  | Time | 0.27 | 2.11 | **0.03** | -0.28 | -1.98 | 0.05 |  |
| **Nitrite Reductase** | (Int) | 3.28 | 10.85 | **<2e-16** | 3.48 | 8.71 | **<2e-16** | (6.23e-04/0.17), (0.05/0.63) |
|  | Time | 0.12 | 1.06 | 0.29 | -0.24 | -2.63 | **0.01** |  |
| **Denitrification** | (Int) | 2.04 | 5.66 | **1.50E-08** | 2.24 | 5.55 | **2.90E-08** | (0.06/0.4) |
|  | Time | -0.03 | -0.24 | 0.81 | -0.31 | -2.85 | **0.004** |  |
| **Nitrite oxireductase** | (Int) | 0.41 | 0.4 | 0.69 | 1.14 | 2.28 | **0.02** | (0.013/0.7), (0.15/0.54) |
|  | Time | -0.23 | -0.98 | 0.33 | -0.57 | -3.14 | **0.002** |  |
| **Nitrification** | (Int) | 1.24 | 1.88 | 0.06 | 0.57 | 2.13 | **0.03** | (0.26/0.71), (0.02/0.12) |
|  | Time | 1.11 | 2.04 | **0.04** | -0.2 | -0.82 | 0.41 |  |
| **Fixation** | (Int) | -15.61 | -1.68 | 0.09 | -1.17 | -1.12 | 0.27 | (0.61/0.99), (0.0006/0.72) |
|  | Time | -9.23 | -1.15 | 0.25 | 0.06 | 0.08 | 0.94 |  |

**Table S8.** Generalized negative binomial regression for the effects of time (days) on all the different urea (organic nitrogen) cycling pathways for burned and unburned plots independently. Significance based on p<0.05, denoted in bold. Negative estimate and z-value scores indicate that time leads to a decrease in the inorganic nitrogen cycling pathways.

|  | **Burned** | | | | **Unburned** | | |  |
| --- | --- | --- | --- | --- | --- | --- | --- | --- |
| **urea cycling** |  | Est. | **z value** | **P value** | **Est.** | **z value** | **P value** | **R2 (B),(Un)** |
| **urea** | (Int) |  | 25.30 | **<2e-16** |  | 30.65 | **<2e-16** | (0.10/0.99) |
|  | Time |  | 1.04 | 0.30 |  | -55.53 | **<2e-16** |  |
| **ornithine biosynthesis** | (Int) |  | 3.16 | **0.002** |  | 2.86 | **0.004** | (0.14/0.4), (0.02/0.02) |
|  | Time |  | -1.55 | 0.12 |  | 0.41 | 0.68 |  |
